# Photosystem II monomeric antenna CP26 has a key role in Non-Photochemical Quenching in *Chlamydomonas reinhardtii*

**DOI:** 10.1101/2022.09.01.506249

**Authors:** Stefano Cazzaniga, Minjae Kim, Matteo Pivato, Federico Perozeni, Samim Sardar, Cosimo D’Andrea, EonSeon Jin, Matteo Ballottari

## Abstract

- Thermal dissipation of the excitation energy harvested in excess, named non-photochemical quenching (NPQ), is one of the main photoprotective mechanisms evolved in oxygenic photosynthetic organisms. Here, the specific function in photoprotection and light harvesting of the monomeric Photosystem II antenna CP26, was investigated in *Chlamydomonas*, model organism for green algae
- CRISPR/Cas9 genome editing and complementation strategies were applied to generate new *cp26* knock-out mutants (named *k6#*) that differently from previous findings, did not negatively affected CP29 accumulation, allowing to compare mutants specifically deprived of CP26, CP29 or both
- The absence of CP26 partially affected Photosystem II activity causing a reduced growth at low or medium light but not at high irradiances. However, the main phenotype observed in k6# mutants was a more than 70% reduction of NPQ compared to wild-type. This NPQ phenotype could be fully rescued by genetic complementation demonstrating that ∼50% of CP26 content compared to wild-type was sufficient to restore the NPQ capacity.
- Our findings demonstrate a pivotal role for CP26 in NPQ induction while CP29 has a crucial function for Photosystem II activity. The genetic engineering of these two proteins could be a promising strategy to regulate photosynthetic efficiency of microalgae under different light regimes.

## INTRODUCTION

Photosynthetic complexes absorb photons and transfer electrons to the redox reactions, leading to the synthesis of NADPH and ATP used to fix CO_2_ into organic molecules (Hill & Scarisbrick, 1940). The two photosystems are organized into two major moieties: reaction center (RC) and antenna subunits (Suga *et al*., 2015). Light-harvesting antenna proteins allow to increase the rate of light energy capturing, efficiently transferring the excitation energy to the RC (Croce & Van Amerongen, 2011). In limiting light conditions, this increased energy capturing is crucial to maximize ATP and NADPH production, required for CO_2_ fixation. However, increased light harvesting capacity may become a potential source of harmful effects upon exposure to high light (Long *et al*., 1994). An excessive light irradiance causes the saturation of the electron transport chain and overload of RCs, blocking or limiting the forward electron transport and preventing the photochemical use of the received energy. Limitation at the acceptor side can extend the lifetime of excited Chls, increasing the probability of generating their triplet states. Triplets, in turn, can react with oxygen, causing the formation of reactive oxygen species (ROS) and leading to photodamage (Neidhardt *et al*., 1998). PSII RC components are particularly sensitive to photodamage (Vass *et al*., 1992) with a specific repair cycle allowing to replace the damaged RC in the membrane (Aro *et al*., 1993). Photodamage can be related to excessive light, but also to variable environmental conditions, that could cause sudden changes in light irradiation (sun flecks), rapidly changing the number of photons received and transiently overload the RCs (Graham *et al*., 2017). Moreover, possible overload of the photosynthetic apparatus can be a consequence of reduced regeneration of ADP and NADP^+^ due to some metabolic impairment or nutrient deficiency. Therefore, an efficient ability to regulate the energetic fluxes along the photosynthetic electron transport chain is a crucial requisite for the fitness of photosynthetic organisms. As a result, photosynthetic organisms need to constantly regulate the mechanisms of light capturing and utilization, to maintain sufficient photochemical activity but avoiding photodamage (Erickson *et al*., 2015).

Evolution provides photosynthetic organisms with mechanisms to protect photosystems and preserve their functionality. Among these, the most responsive is the non-photochemical quenching (NPQ) of Chls, that rapidly dissipates the excess energy absorbed as heat, preventing triplet states formation (Horton, 1996). NPQ response is triggered through a feedback mechanism by thylakoid lumen acidification, which is caused by saturation of the photosynthetic electron transport chain (Genty *et al*., 1989). Understanding the molecular mechanism of NPQ activation and deactivation process have a great biotechnological relevance to improve the fitness and yield of photosynthetic organisms (Kromdijk *et al*., 2016; Głowacka *et al*., 2018). Microalgae raised interest for biotechnological applications, being photosynthetic microorganisms which can be cultivated in industrial environment as biofactories for bioproducts of interest, like food additives, nutraceuticals, antioxidants, biostimulants, and even biofuels (Medipally *et al*., 2015; Bernaerts *et al*., 2019; Camacho *et al*., 2019; Koyande *et al*., 2019; Rani *et al*., 2021). In the model organism for green algae, *Chlamydomonas reinhardtii*, the key proteins responsible for this mechanism activation are the pigment binding proteins LHCSR1 (Light Harvesting Complex Stress Related 1) and LHCSR3 subunits, with the latter having a predominant role (Peers *et al*., 2009). Expression of LHCSRs is strongly induced upon high light exposure. LHCSR proteins were reported to sense lumen acidification upon protonation of specific exposed residues, thus being able to switch from a light-harvesting state to a quenched one triggering NPQ (Peers *et al*., 2009; Liguori *et al*., 2013; Ballottari *et al*., 2016; Troiano *et al*., 2021).

While LHCSR is the trigger, it needs to interact with other antenna proteins of the photosystems to efficiently induce NPQ (Tokutsu & Minagawa, 2013; Dinc *et al*., 2016). In *C. reinhardtii*, PSII light antenna are organized in two different moieties: trimeric major light-harvesting complexes (LHCII) and monomeric subunits (Shen *et al*., 2019). Trimeric LHCII are encoded by nine genes called *lhcbm1*–*lhcbm9* (with M referring to ‘major’ antenna complex) (Minagawa & Takahashi, 2004; Drop *et al*., 2014). Conversely, CP26 and CP29 are the two the monomeric LHC subunits which have a key role as linker between the RC and the external LHCII (Drop *et al*., 2014; Semchonok *et al*., 2017; Cazzaniga *et al*., 2020). Indeed *C. reinhardtii* PSII homodimer RC is surrounded by two antenna layers (Tokutsu *et al*., 2012): the inner layer is composed of CP29, CP26, and S-LHCII (S, strongly bound), forming with the core the C_2_S_2_ particle, whereas the outer layer is made of M-LHCIIs (moderately bound) forming with C_2_S_2_, the C_2_S_2_M_2_ supercomplex. An additional LHCII (N-LHCII) can be connected directly to the PSII core in the position that in vascular plants is occupied by the third monomeric antenna CP24 (absent in *C. reinhardtii*), forming the larger C_2_S_2_M_2_N_2_ (Drop *et al*., 2014).

The different LHC subunits have similar biochemical and biophysical properties (Natali & Croce, 2015), but despite these structural similarities, different antennas have been suggested to have different roles in the light harvesting mechanism and photoprotection. Indeed, the NPQ phenotype of specific *lhcbm* knock out or knock down mutant led to the suggestion of peculiar roles in NPQ for LHCBM1 and LHCBM4-6-8. In the case of monomeric antenna LHC subunits, single mutant on CP26 and CP29 genes, called k6 and k9 respectively, and a double CP26 and CP29 mutant, called *k69*, were recently reported in *C. reinhardtii* showing a strong reduction of light harvesting and NPQ mechanisms in the absence of both antenna proteins (Cazzaniga *et al*., 2020). However, in the *k6* mutant previously described the absence of CP26 also caused the absence of CP29 resulting in a similar phenotype of *k6* and *k69* mutants. Detailed information on the specific role of CP26 and CP29 in photoprotection and light harvesting are thus still missing. In this study, we further characterize the role of monomeric antenna in *Chlamydomonas reinhardtii* using CRISPR/Cas9 genome editing and complementation strategies in *C. reinhardtii* obtaining new *cp26* mutants (herein named *k6#*) where CP29 expression was similar to the Wt case. The new *k6#* mutants were characterized by partially reduced photosynthetic efficiency, reduced growth compared to its parental strain, and a strong impairment in NPQ induction. This finding, supported by genetic complementation, demonstrates an important difference between the function of CP26 in green algae and vascular plants: in the latter indeed the absence of CP26 had a minor effect, if any on NPQ induction (de Bianchi *et al*., 2008).

## MATERIALS AND METHODS

### Culture conditions

*C. reinhardtii* CC503 (wild-type, Wt) and mutant strains were grown at 24°C in high-salts (HS) medium (Kropat *et al*., 2011) on a rotary shaker in Erlenmeyer flasks under continuous illumination with white LED. Cell densities were measured using Countess II FL Automated Cell Counter (Thermo Fisher Scientific). Single mutants on CP29 (*k9*) and double mutant on CP26 and CP29 (*k69*) were obtained from (Cazzaniga *et al*., 2020).

### Generation of C. reinhardtii mutants by CRISPR-Cas9 genome editing

To generate knock-out CP26 mutants, CRISPR-Cas9 RNP complex-mediated genome editing method was performed according to (Kim, E *et al*., 2020) and (Joo *et al*., 2022) (see Supporting Information Methods S1). To verify the integration and copy number of the aphVII gene in the genome, the genomic DNA was digested by *Kpn*I or *Nco*I, and Southern blot analysis was performed using aphVII gene probe for detection of DNA integration. Bands were visualized using the Gene Images AlkPhos Direct Labeling and Detection System (Amersham, Little Chalfont, Bucks, United Kingdom).

### Mutant complementation

Vector used for CP26 knockout mutant complementation was obtained by fusion PCR merging the 1000bp upstream 5’UTR of CP29 gene (Cre17.g720250) with the whole CP26 gene (Cre16.g673650) deprived of promoter region and extended of 300bp downstream 3’UTR. In the case of CP29 complementation in *k69* background the genomic sequence of CP29 gene from 1000bp upstream of 5’ UTR to 300bp downstream of 3’ UTR was used. These sequences were cloned into pOpt2 vector system and used for transformation. Nuclear transformation was carried out by glass bead as previously described (Perozeni, F *et al*., 2020) using 10 μg of linearized plasmid followed by selection of transformants on TAP agar plates (Kropat *et al*., 2011) supplied by paromomycin (10 mg/L) for 6–7 days.

### Membrane preparation, Gel electrophoresis and immunoblotting

Thylakoid membranes were isolated as previously described (Bonente *et al*., 2008). SDS-PAGE analysis was performed using the Tris-Tricine buffer system (Schägger & von Jagow, 1987) followed by Coomassie blue staining. For immunotitration, thylakoid samples were loaded for each sample and electroblotted on nitrocellulose membranes; then, proteins were quantified with an alkaline phosphatase–conjugated antibody system. αCP26 (AS09 407), αCP29 (AS04 045), αLHCII (AS01 003) αCP43 (AS11 1787), αPSAA (AS06 172), αLHCSR1 (AS14 2819), αLHCSR3 (AS14 2766) antibodies were purchased from Agrisera (Sweden). Non-denaturing Deriphat-PAGE was performed following the method developed in (Peter *et al*., 1991). Thylakoids concentrated at 1 mg/ml Chls were solubilized with a final 0.8% α-DM, and 30 μg of Chls were loaded in each lane.

### Pigment and spectroscopy analysis

Pigments were extracted from intact cells using 80% acetone buffered with Na_2_CO_3_ and analyzed by reverse-phase HPLC (Lagarde *et al*., 2000) as described in (Perozeni, F. *et al*., 2020). 77 K fluorescence emission spectra on frozen cell were registered using a BeamBio custom device equipped with USB2000 Ocean Optics spectrometer (Ocean Optics).

### Time-resolved fluorescence on whole cells

Time-resolved fluorescence measurements were carried out using a Ti: sapphire laser (Chameleon Ultra II, Coherent, repetition rate of 80 MHz and 140 fs pulse width). A β-barium borate (BBO) crystal was used to frequency double the 950 nm laser output in order to generate the 475 nm excitation wavelength. The fluorescence signal from the cells was focused on the entrance slit of a spectrograph (Acton SP2300i, Princeton Instrument). A streak camera (C5680, Hamamatsu), equipped with the Synchroscan sweep module, which provides spectral-temporal matrices with spectral and temporal resolutions of ∼1 nm and ∼20 ps, respectively, for the temporal window of 2 ns. A CCD (Hamamatsu ORCA-R2 C10600) recorded the streak image. The measurements were carried out in cuvettes with optical path-length of 10 mm and the sample was under constant circulation from a reservoir of 30 mL sample volume through a tube and peristaltic pump to avoid any photodamage of the samples.

### Measurements of photosynthetic activity

Photosynthetic parameters ΦPSII, qL, electron transport rate (ETR), and NPQ were obtained by measuring with a DUAL-PAM-100 fluorimeter (Heinz–Walz) chlorophyll fluorescence of intact cells, at room temperature in a 1 × 1-cm cuvette mixed by magnetic stirring. ΦPSII, qL, and ETR were measured and calculated according to (Baker, 2008) and (Van Kooten & Snel, 1990) upon 20 min of illumination. NPQ measurements were performed on dark-adapted intact cells as described in (Cazzaniga *et al*., 2020) with the following modifications: cells were adapted to high light only for two days before NPQ measurements and far-red light exposure was performed for 2 minutes before turning on the actinic light. PSII functional antenna size was measured from fast chlorophyll induction kinetics induced with a red light of 11-μmol photons m^−2^ s^−1^ on dark-adapted cells (∼2·10^6^ cells/ml) incubated with 50-μM DCMU. The reciprocal of time corresponding to two thirds of the fluorescence rise (τ2/3) was taken as a measure of the PSII functional antenna size (Malkin *et al*., 1981). The oxygen evolution activity of the cultures was measured at 25°C with a Clark-type O2 electrode (Hansatech), as described previously (Cazzaniga *et al*., 2020). NPQ acid induction in the dark was induced as previously described (Tian *et al*., 2019). See Supporting Information Methods S1 for additional details.

## RESULTS

### Expression of CP29 protein in absence of CP26 and generation of CP26 deficient mutant in Chlamydomonas

The absence of CP29 in the previously reported *k6* mutant suggested a possible destabilization of CP29 binding to PSII in absence of CP26 (Cazzaniga *et al*., 2020). To test this hypothesis, CP29 complementation was attempted in the double mutant *k69* background introducing the whole CP29 genomic sequence including, native promoter, 5’-UTR, 3’-UTR and terminator (Supporting Information Fig. S1). Transformant lines successfully accumulated CP29 subunit to a similar level compared to Wt, even in the absence of CP26. Despite the CP29 accumulation, the new complemented strains were characterized by a strong reduction of NPQ similar to their background (*k69* mutant), suggesting a pivotal role for CP26 in NPQ. These finding prompted us to generate new CP26 mutants to investigate the role of this antenna protein in *C. reinhardtii*. A reverse genetic approach was applied to knock out the CP26 gene (lhcb5, Cre16.g673650) in a Wt background through CRISPR/Cas9-mediated gene disruption (Baek *et al*., 2016; Kim, J *et al*., 2020) (Fig. 1). The sgRNA target located in exon 4 yielded successful results, leading to non-homologous end joining and Cas9-mediated hygromycin-resistance (HygR; aphVII) gene cassette insertion in the target site (Fig. 1a). Insertion of HygR gene cassette in the coding DNA sequence (CDS) of the target gene was confirmed by PCR and Sanger sequencing (Fig.1, Supporting Information Fig. S2 and Table S1) and the lines were named *k6#1-11*. Ten mutants were validated by immunoblot analysis with a specific antibody against CP26 confirming the absence of the protein (Fig. 1d, Supporting Information Fig. S1b). Southern blot analysis on the selected lines showed that *k6#4* and *k6#8* contained a single insertion of the HygR gene in the genome, while the others displayed multiple insertions (Supporting Information Fig. S1c). These two mutants were considered for subsequent analysis.

**Figure 1.**
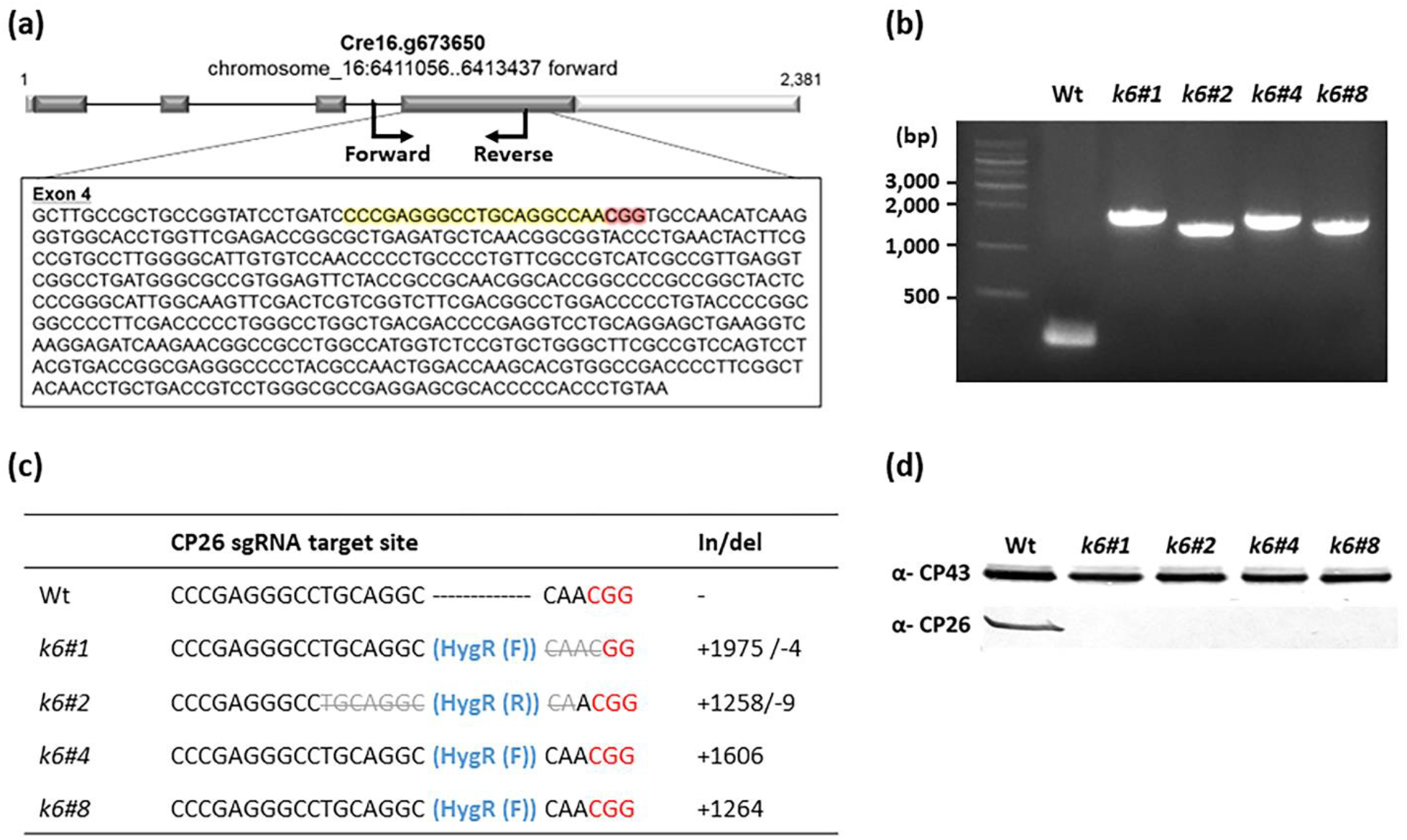
CP26 knock-out by CRISPR/Cas9 genome editing. (a) Location of the Cas9 target site in the CP26 gene. Light gray colored- and dark gray colored-box indicate UTR regions and exons, respectively. The nucleotides highlighted in yellow and red indicate the sgRNA sequence and its’ PAM sequence, respectively. Black arrows indicate the primer sets for colony PCR (b) Colony PCR of the selected CP26 knock-out mutants using the specific primers designed near the Cas9 target site of CP26. (c) Sanger sequencing of the selected CP26 knock-out mutants. In/del values represent inserted or deleted nucleotides in sgRNA target site of CP26 in wild-type (Wt). Hygromycin-resistance gene (HygR/ aphVII) cassette inserted in the target cited of CP26 gene was identified with either forward (F) or reverse (R) orientation. The size of the HygR cassette is 1,624 bp. The deleted sequence in the sgRNA target site was highlighted by a gray-colored line (d) Immunoblot analysis of the four selected CP26 knock-out mutants with a specific antibody directed against CP26. Immunoblot against CP43 was added as loading control

### Pigment content and Pigment-Protein Complexes stoichiometry in the mutant without CP26

Pigment content was analyzed in Wt and *k6#* cell cultures grown in photoautotrophy at 80-100 μmol m^−2^ s^−1^ for at least three generations (Table 1). The mutant, compared to its parental line, displayed a similar Chl content per cell, but a slightly lower Chl a/b ratio. The Chl per carotenoids ratio and the carotenoid composition were comparable between the two genotypes, as confirmed by the high-performance liquid chromatography (HPLC) analysis showing only minor and not significant differences in loroxanthin and antheraxanthin content. At this light intensity, zeaxanthin accumulation was not detected.

**Table 1.** Pigment analysis and Fv/Fm. Data were collected from cells grown in photoautotrophy at 80-100 μmol photons m-2 s-1. Single carotenoids values were normalized to 100 chlorophylls. In the case of *k6#* the results obtained were obtained from two independent mutant lines. Data are expressed as mean ± SD. (n = 4). Values that are significantly different (Student’s t-test, P < 0.05) from the wild-type (Wt) are marked with an asterisk (*). Abbreviations: Chl, chlorophyll; Car, carotenoids; Neo, neoxanthin; Lor, loroxanthin; viola, violaxanthin; Ant, anteraxanthin; Lut, lutein; β-Car, β-Carotene.

|  |  |  |  |  | Car/100 Chl |  |  |  |  |  |
| --- | --- | --- | --- | --- | --- | --- | --- | --- | --- | --- |
| | Chl<br>(pg)/cel<br>l | Chl<br>a/b | Chl/Car | Fv/F<br>m | Neo | Lor | Viol | Ant | lut | β-Car |
| Wt | 1.40 ±<br>0.07 | 2.65<br>±<br>0.04 | 3.16 ±<br>0.07 | 0.72 ±<br>0.01 | 3.43 ±<br>0.59 | 5.12 ±<br>0.36 | 3.30 ±<br>0.08 | 0.39 ±<br>0.04 | 13.85 ±<br>0.50 | 4.42 ±<br>0.71 |
| *k6#* | 1.47 ±<br>0.22 | 2.48<br>±<br>0.04 | 3.16 ±<br>0.03 | 0.67 ±<br>0.02* | 3.60 ±<br>0.96 | 3.99 ±<br>0.37 | 3.23 ±<br>0.27 | 0.15 ±<br>0.14 | 15.03 ±<br>0.90 | 4.21 ±<br>0.47 |

A partial rearrangement of the photosynthetic complexes in *k6#* lines could account for a lower Chl a/b ratio, being Chl b only bound by antenna subunits while Chl a is present in all pigment-binding complexes. To assess this hypothesis, we measured the relative amounts of PSI, PSII, LHCII and CP29 using specific antibodies against the different photosynthetic subunits (Fig. 2). In particular, CP43 and PsaA were used as proxies for PSI and PSII content, respectively, while LHCII content was investigated using an antibody recognizing the different LHCBM subunits of *C. reinhardtii* (Girolomoni *et al*., 2017). Compared to Wt, the *k6#* mutant showed increased PSI/PSII and LHCII/PSII ratios. In particular, LHCII content per PSII was almost doubled, suggesting a possible compensation of the absence of monomeric antenna by an increase of the LHCII protein level (Fig. 2a). In the case of CP29, the CP29/PSII ratio was similar in *k6#* compared to Wt, as in the case of *k69* mutant complemented with CP29 (Supporting Information Fig. S1). The organization of pigment binding complexes was further analysed by Non-denaturing Deriphat-PAGE, upon solubilisation of thylakoid membranes with dodecyl-α-D-maltoside (α-DM) (Fig. 2c). Different pigment binding bands were resolved: *k6#* mutant showed a relative increase of the bands corresponding to antenna proteins and to PSI compared to PSII core. It is important to note that the reduced Chl a/b ratio and the increased PSI/PSII and LHCII/PSII ratios observed in *k6#* are essentially in line with previous observations in the *C. reinhardtii* mutant *k9*, where CP29 was absent, while in the double mutant *k69* the LHCII and PSI per PSII content were further increased (Cazzaniga *et al*., 2020). However, it is important to point out that while the CP29/PSII was similar in *k6#* and Wt, in the case of *k9* mutant a ∼50% increase in CP26/PSII ratio compared to the Wt case was observed (Supporting Information Fig. S3), likely to partially compensate for the absence of CP29.

**Figure 2.**
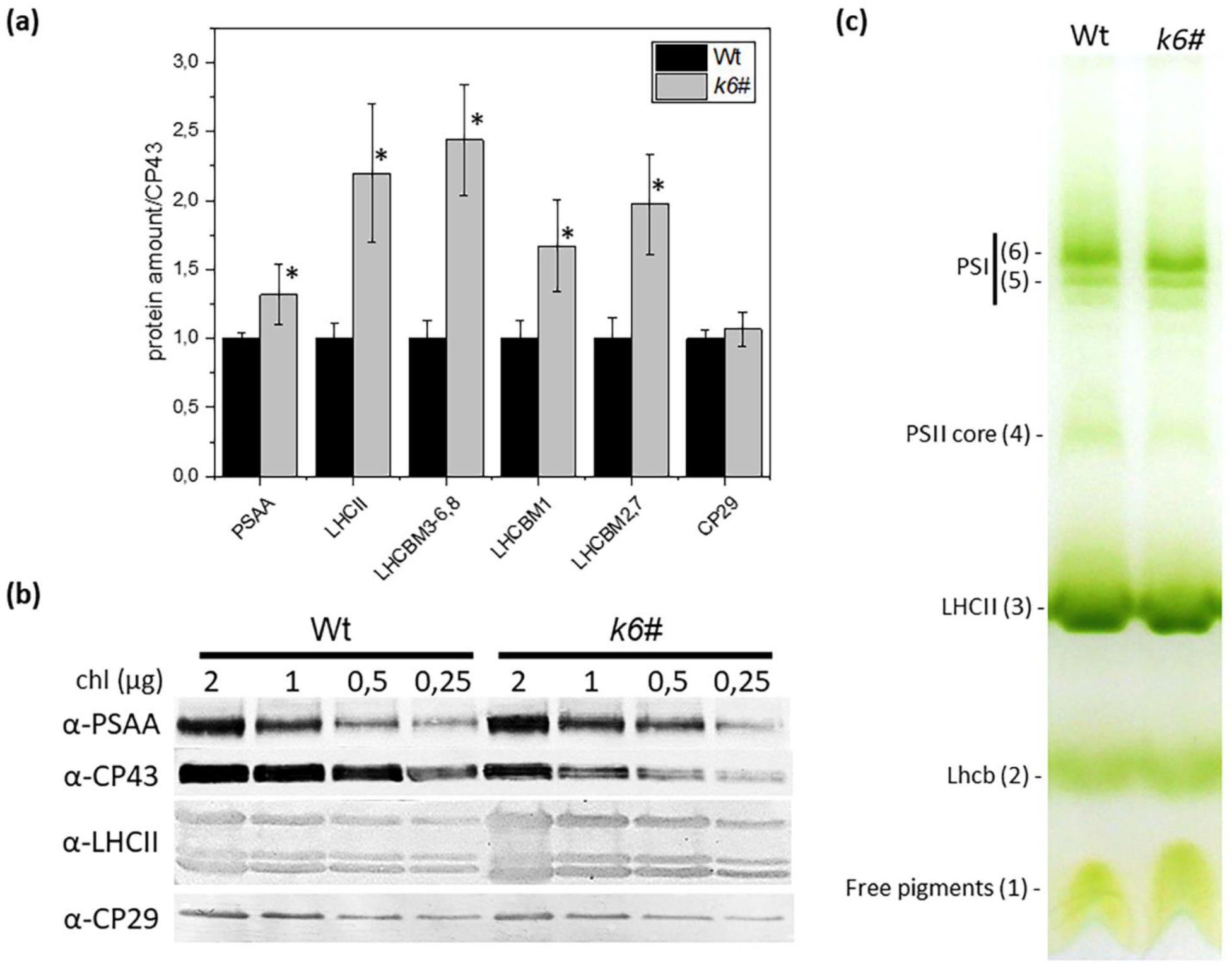
Organization of the photosynthetic complexes. (a) Immunotitration of wild-type (Wt) and *k6#* thylakoid proteins using specific antibodies against PSI core (PSAA), LHCII and CP29. Data were corrected for CP43 amount and normalized to Wt ratio. LHCII is shown as total amount (LHCII) or divided for three bands that are separated on the Tris-Tricine buffer system: the upper one corresponding to *lhcbm3-6* and *lhcbm8* gene products, the one in the middle to *lhcbm1* and the lower one to *lhcbm2* and lhcbm7 (Girolomoni et al., 2017). Data are expressed as mean ± SD. (n = 4). *k6#* values that are significantly different (Student’s t-test, P < 0.05) from the Wt are marked with an asterisk (*). (b) Image of western blot used for the Immunotitration. Specific antibodies were used on cellulose lanes loaded with 2, 1, 0.5 and 0.25 μg of Chls. (c) Thylakoid pigment-binding complexes were separated by non-denaturing Deriphat-PAGE upon solubilization with 0.8% α-dodecyl-maltoside. Thylakoids corresponding to 30 μg of Chls were loaded in each lane. Composition of each band is indicated.

### Light harvesting efficiency

The consequences of CP26 absence on the activity of PSII were initially evaluated by measuring PSII maximum quantum efficiency (F_v_/F_m_) in dark-adapted cells (Butler, 1973), revealing a significant decrease in the mutant (0.67 vs 0.72 in *k6#* and Wt, respectively) (Table 1). Such difference is attributable to a higher basal fluorescence (F_0_) in the mutant when exposed to weak measuring light, as reported by the fluorescence values normalized to the Chl amount (Supporting Information Fig. S4): a higher F_0_ value in the *k6#* indicates that a fraction of the absorbed light energy is not efficiently directed toward photochemistry but re-emitted as fluorescence.

This alteration can be observed also by measuring fluorescence emission spectra at 77K (Fig. 3a): *k6#* cells were characterized by an additional shoulder at ∼680 nm, usually assigned to “disconnected” LHC complexes, meaning LHC proteins are impaired in energy transfer to Photosystems, thus re-emitting fluorescence at shorter wavelengths (Garnier *et al*., 1986). *k6#* mutant was thus characterized by an increased LHCII content, which however are partially not able to transfer energy to PSII: a similar phenotype could be observed in absence of CP29 (*k9* mutant) or, to a higher extent, in the double *k69* mutant, without both CP26 and CP29 (Supporting Information Fig. S5a), consistently with previous data (Cazzaniga *et al*., 2020). The light harvesting capacity of PSII, or functional PSII antenna size, was thus evaluated from the kinetics of Chl a fluorescence induction in whole cells in limiting light conditions, upon treatment with PSII inhibitor DCMU: in this condition the rise time of chlorophyll fluorescence is inversely proportional to PSII light harvesting feature (Malkin *et al*., 1981). *k6#* showed a functional antenna size similar to the Wt case, even with a doubled LHCII/PSII ratio, further suggesting a reduced antenna-to-core energy transfer efficiency in this mutant (Fig. 3b). Energy transfer to PSII RC was thus investigated on whole cells by measuring time-resolved fluorescence decay on Wt and *k6#* cells using 475-nm excitation (Fig. 3c). In absence of CP26, the fluorescence decay traces were slower compared to the Wt case. Similar results could be obtained in the case of *k9* and *k69* (Supporting Information Fig. S5b). As reported in Supporting Information Table S2, fitting analysis with exponential functions allow to estimate the average lifetimes of fluorescence decay, which were increased in *k6, k9* and *k69* compared to Wt, with the longest lifetime in the double mutant. These results confirm that in absence of CP26 energy transfer from LHCII to RC was partially impaired, with a more evident consequences when both CP26 and CP29 were absent.

**Figure 3.**
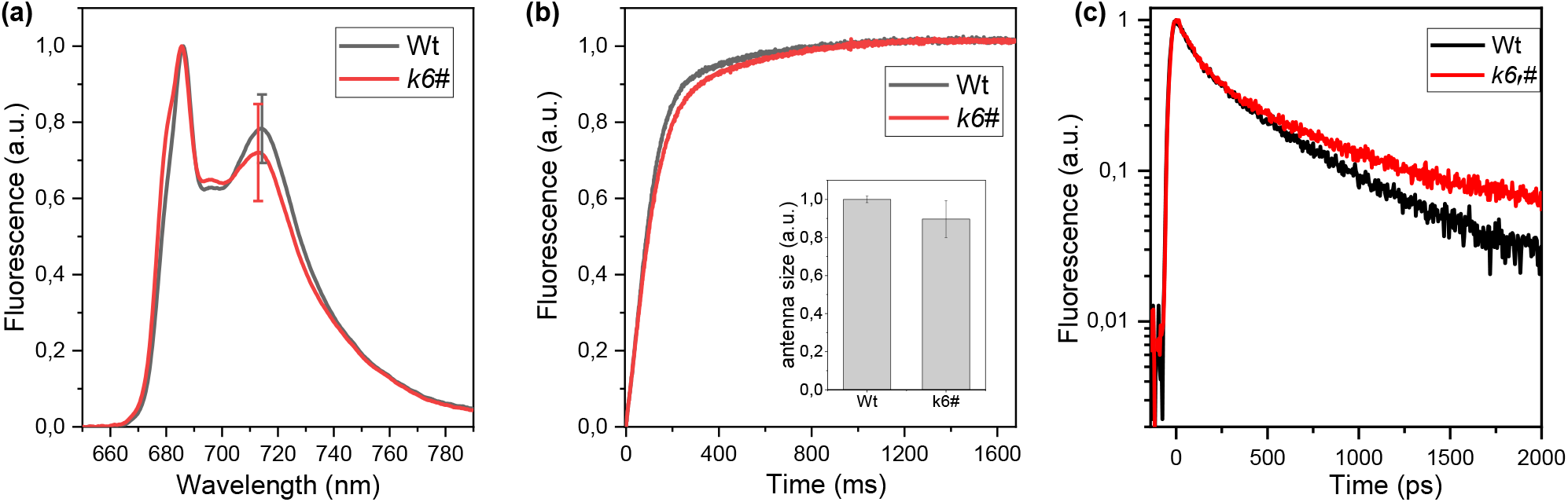
Light harvesting. (a) Low-temperature fluorescence emission spectra of wild-type (Wt, black) and *k6#* (red) cells excited at 475 nm. Emission spectra are normalized at the maximum emission on PSII peak. Error bars are reported as standard deviation (n = 4) indicated on the PSI peak. In the case of *k6#* mutant the results reported were obtained from two independent mutant lines. (b) Variable Chl fluorescence was induced with a weak red light on dark-adapted cells (2 · 10^6^ cells/ml) in presence of 50 μM of DCMU. The trace for Wt and *k6#* are the average of 12 curve from three different experiments. The reciprocal of time corresponding to two-thirds of the fluorescence rise (1/τ2/3) is as a measure of the PSII functional antenna size and it is shown in the inset normalized to the Wt. Error bars are reported as standard deviation (n = 12). In the case of *k6#* mutant the results reported from two independent mutant lines. (c) Time-resolved fluorescence decays of Wt and *k6#* whole cells. In the case of *k6#* mutant the results reported in a, b and c were obtained from two independent mutant lines.

PSII photochemical activity was then investigated in *k6#* and its parental strain by measuring PSII operating efficiency (ΦPSII), PSII electron transport rate (ETR) and photochemical quenching (1-qL) at different light intensities (Baker, 2008). As reported in Fig. 4, these photosynthetic parameters were not significantly different between the Wt and *k6#* mutant. The only exception is represented by the ΦPSII at low light intensities which was lower in *k6#* mutants. These results are different compared to the *k9* and *k69* case, where a strong reduction in ETR and ΦPSII was observed, especially in the double *k69* mutant (Supporting Information Fig. S5), consistent with previous findings (Cazzaniga *et al*., 2020). The influence of CP26 on light dependent oxygen production was then investigated at different actinic lights (Fig. 4d). Despite having a similar maximum light dependent oxygen production rate (Pmax), *k6#* mutant was characterized by a reduced oxygen production at low and intermediate irradiances (Fig. 4d, Supporting Information Table S3). The respiration in the dark was not affect by CP26 absence (Supporting Information Table S3). However, a more evident decrease in oxygen production could be observed in the absence of CP29 in *k9* mutant and even further in the double mutant *k69*. From these data, the *k6#* showed a lower photochemical efficiency compared to Wt only at low-medium light, while a more severe reduction of PSII activity, even at high irradiances, could be observed in the absence of CP29 or both CP26 and CP29 (Supporting Information Fig. S5).

**Figure 4.**
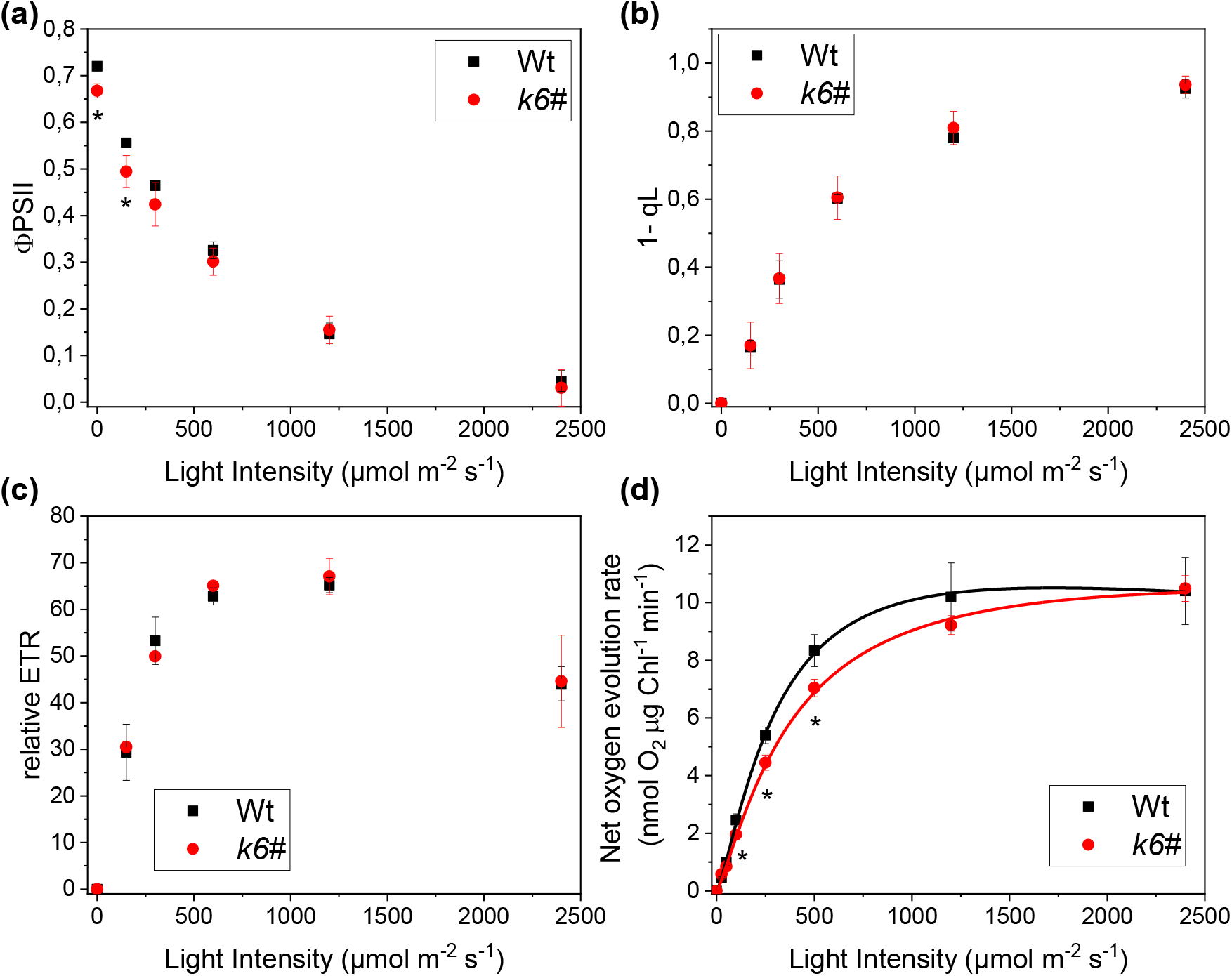
Photosynthetic electron flow. Dependence of the (a) PSII operating efficiency (ΦPSII), (b) relative electron transport rate (ETR), (c) 1-qL (reduced QA fraction), and (d) photosynthetic O2 evolution on actinic light intensity for wild type (Wt, black) and *k6#* (red). Net photosynthetic rate data were fitted with Hill equation. In the case of *k6#* mutant the results reported were obtained from two independent mutant lines. Error bard are reported as standard deviation (n > 3). *k6#* values that are significantly different (Student’s t-test, P < 0.05) from Wt are marked with an asterisk (*).

### Biomass productivity

To evaluate how these defects in *k6#* could affect biomass yield, growth curves of both Wt and *k6#* strains were monitored in photoautotrophy at low (40 μmol m^−2^ s^−1^), medium (80-100 μmol m^−2^ s^−1^) and high light (700-800 μmol m^−2^ s^−1^) (Fig. 5). Under low or medium irradiances, the absence of CP26 caused a significant delay in growth rate reaching in the case of *k6#* mutant a cell density around half of the Wt. However, in the absence of CP29 (*k9* mutant) or both CP26 and CP29 (*k69* mutant) a much stronger reduction in biomass productivity could be observed (Supporting Information Fig. S6). At higher light intensities, where light is not limiting, the growth curves of Wt, *k6, k9* and *k69* strains were similar (Fig. 5c, Supporting Information Fig. S6).

**Figure 5.**
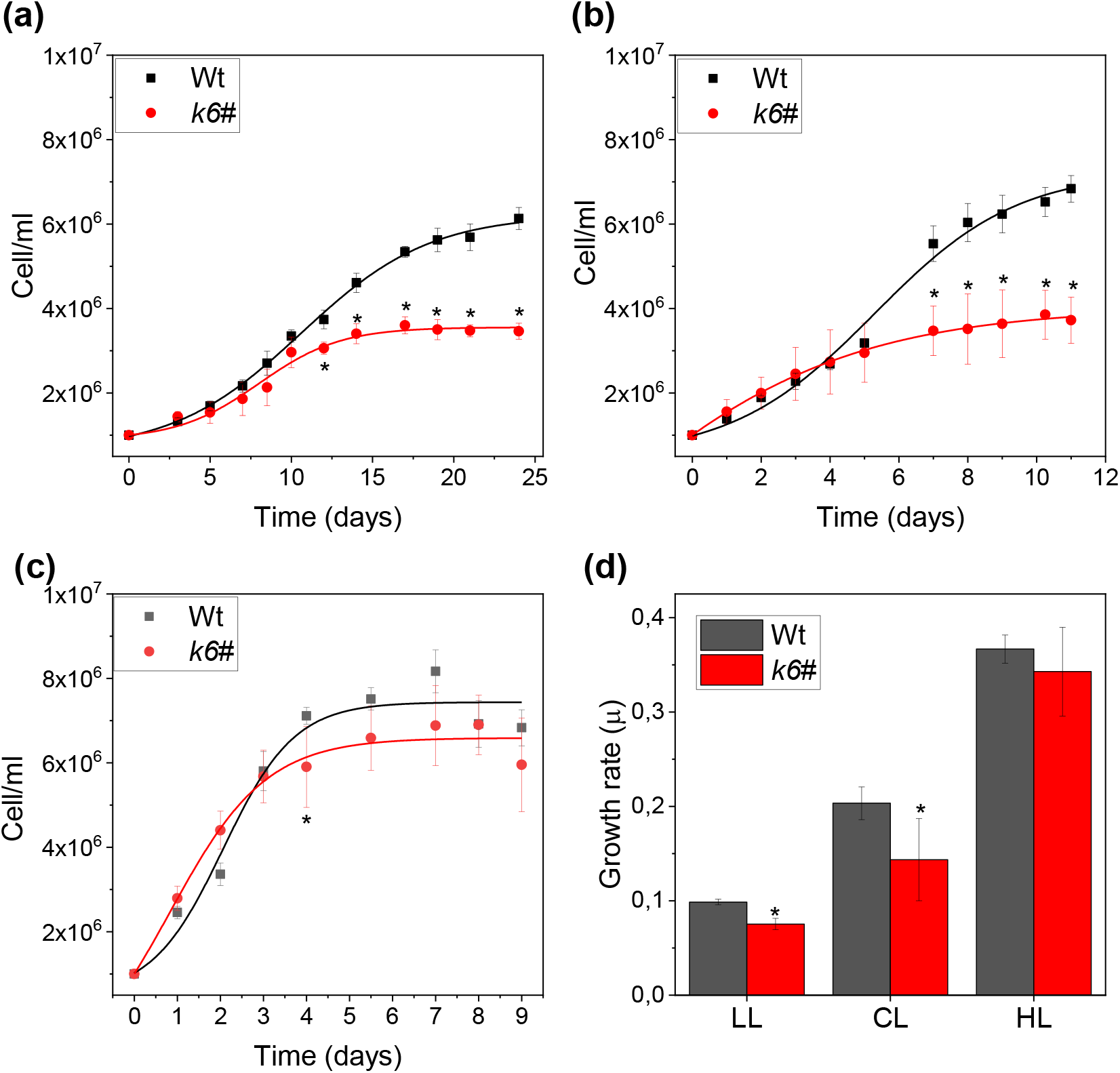
Growth curves under autotrophic conditions. Wild-type (Wt, black) and *k6#* (red) where grown at (a) low (30-40 μmol m^−2^ s^−1^,LL), (b) control (100-120 μmol m^−2^ s^−1^,CL) or (c) high light (700-800 μmol m^−2^ s^−1^,HL) and the cell content was measured. Growth curves were fitted with Hill equation. (d) specific growth rate (μ) calculated for all the different light conditions. Error bars are reported as standard deviation (n > 3). *k6#* values that are significantly different (Student’s t-test, P < 0.05) from Wt are marked with an asterisk (*). In the case of *k6#* mutant the results reported were obtained from two independent mutant lines.

### Non-Photochemical Quenching induction

Previous work demonstrated that the photoprotective induction of NPQ was essentially impaired in the absence of CP26 and CP29, but still active in the absence of CP29 only (Cazzaniga *et al*., 2020). The availability of a single *k6#* mutant depleted of CP26 allows to investigate the specific function of this LHC complex in NPQ induction in *C. reinhardtii*. Because LHCSR proteins are required to induce NPQ (Peers *et al*., 2009), Wt and *k6#* strains were exposed to high light (1200 μmol m^-2^ s^-1^) for two days to induce LHCSR1 and LHCSR3 (Allorent *et al*., 2013). Immunoblotting analysis revealed an increased content per PSII of both LHCSR1 and LHCSR3 in *k6#* compared to Wt (Supporting Information Fig. S7). Despite the increased LHCSR content per PSII, NPQ induction in k6# mutant was strongly reduced compared to the Wt case with a residual NPQ ranging from ∼6% at 150 μmol photons m^−2^ s^−1^ to ∼30% at 2400 μmol photons m^−2^ s^−1^ in the mutant (Fig.6, Supporting Information Fig. S8). In the same measuring conditions, the NPQ was further reduced in the *k69* double mutant (Supporting Information Fig. S9), with a residual NPQ compared to Wt ranging from 0% at 150 μmol photons m^−2^ s^−1^ to 10% at 2400 μmol photons m^−2^ s^−1^, consistently with previous findings (Cazzaniga *et al*., 2020). The absence of CP29 instead had a minor effect on NPQ with a residual NPQ compared to the Wt case ranging from 50% at 150 μmol photons m^−2^ s^−1^ to 90% at 2400 μmol photons m^−2^ s^−1^.

NPQ is triggered by thylakoid lumen acidification. The NPQ phenotype observed in *k6#* mutant could in principle be related to a reduced capacity in the mutant to generate a proton gradient across the thylakoid membrane upon illumination. To overcome this possible bias, light independent NPQ activation was induced by direct lumen acidification adding acid solution to the cells as previously reported (Tian *et al*., 2019). Cells were first adapted to high light for two days to induce LHCSR and then maximum fluorescence emissions were measured applying saturating pulses, before (F_m_) and after (F_m_^a^) decreasing the pH with acid solution (Fig. 7). In the Wt, the acidification induced a strong decrease of F_m_^a^ compared to F_m_ with a calculated NPQ of 1.76, whereas in *k6#*, the same procedure induced only a limited decrease of the F_m_^a^ and a NPQ of 0.52. These differences between Wt and *k6#* are comparable to the ones obtained inducing NPQ by light exposure, further suggesting that the impaired quenching in *k6#* is mainly caused by a hampered NPQ mechanism rather than an impairment in the thylakoid lumen acidification. To validate the procedure, we tested direct acidification on null NPQ mutant (*npq4 lhcsr1*, lacking both LHCSR1 and LHCSR3) and its parental strain (4a+), observing an absence of pH dependent quenching in the absence of LHCSR subunits (Supporting Information Fig. S10). The same approach tested on *k9* and *k69* cells showed that the mutant without CP29 had a level of NPQ similar to Wt while the double mutant *k69* showed a strongly reduced level of quenching (Supporting Information Fig. S11).

**Figure 6.**
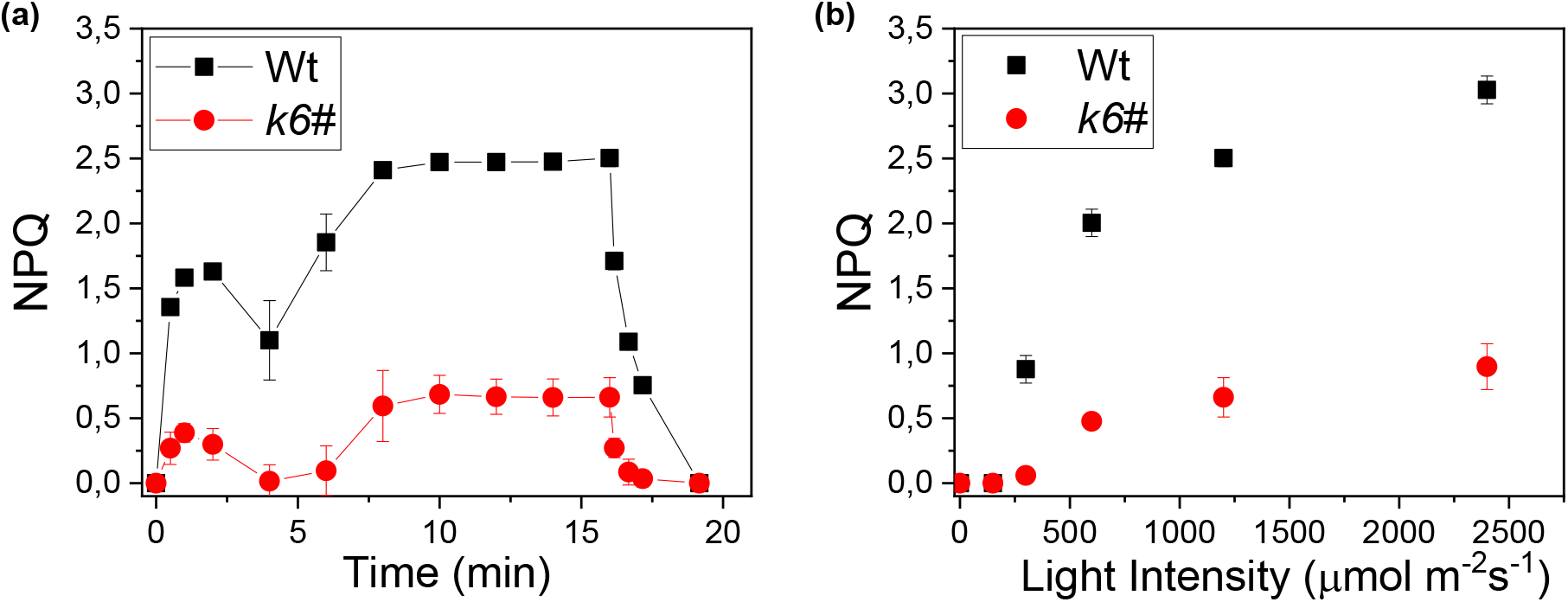
Nonphotochemical quenching. (a) Measurement of NPQ kinetic on wild-type (Wt, black) and *k6#* (red) cells using actinic lights of 1,200 μmol photons m^−2^ s^−1^. (b) NPQ value after 15 min of illumination at different actinic light intensities. Error bars are reported as standard deviation (n >4). In the case of *k6#* mutant the results reported were obtained from two independent mutant lines.

**Figure 7.**
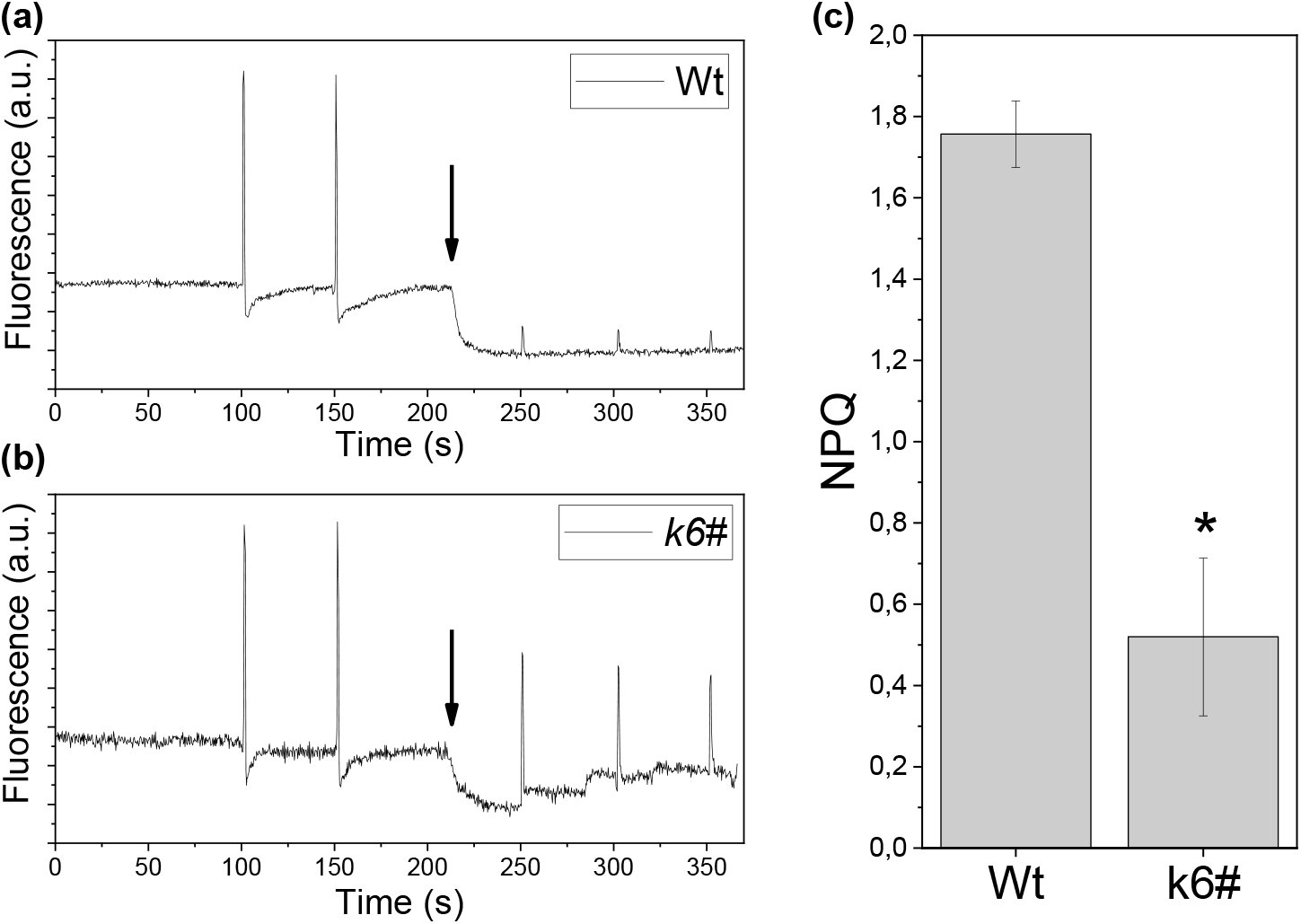
Acid induced quenching. (a,b) PAM Chl fluorometer trace of wild-type (Wt) (a) and *k6#* (b) before and after acidification with acetic acid. The back arrow indicates when the acid was added. Fluorescence was recorded in the dark with two saturating pulses before and two after the acidification. (c) NPQ calculated for each trace as (Fm-Fm^A^)/Fm^A^ where Fm and Fm^A^ are the fluorescence before and after acidification. Error bars are reported as standard deviation (n > 3). *k6#* values that are significantly different (Student’s t-test, P < 0.05) from Wt are marked with an asterisk (*).

### Rescue of NPQ phenotype by genetic complementation

NPQ reduction was the main phenotype observed in *k6#* mutant compared to Wt. Therefore, a genetic complementation approach was performed to correlate this phenotype adequately with the absence of CP26. First, we tried to use the whole CP26 genomic sequence including, native promoter, 5’-UTR, 3’-UTR and terminator (Supporting Information Fig. S12). However, we were unable to synthetize or amplify the 1000 bp upstream the CDS, due to its complexity and to the presence of an un-sequenced region into the genome. To overcome this issue, we decided to exploit the 1000 bp upstream region of the other monomeric antenna CP29 (Cre17.g720250) as putative promoter, which did not pose the same challenge as the CP26 region. This chimeric construct (CP29, 1000 bp upstream-CP26 gene from 5’UTR to 3’UTR-CP26, 300 bp downstream) was then introduced in the *k6#8* mutant line, resulting in the successful accumulation of CP26 protein. The CP26 expressing lines in *k6#* background, were named *k6#c* (*ko cp26* complemented): different levels of CP26 expression were observed in *k6#c* lines, ranging from ∼10% to 200% of the Wt CP26/PSII ratio (Supporting Information Fig. S13). *k6#c* lines were adapted for two days in high light and the NPQ was measured upon exposure to 1200 μmol photons m^−2^ s^−1^ actinic light. As reported in Fig. 8a, in the case of five different complemented lines displaying a CP26/PSII ratio comparable to Wt (Supporting Information Fig. S13), the NPQ induction kinetics were similar between Wt and the selected *k6#c* lines, highlighting that the NPQ phenotype in *k6#* was specifically due to the absence of CP26. Moreover, in these lines the Fv/Fm values were similar to the Wt case, confirming the correlation between the lack of CP26 expression and decreased PSII efficiency (Supporting Information Fig. S13). NPQ values obtained for the different complemented lines expressing different levels of CP26 were then plotted as a function of CP26/PSII ratio (Fig. 8b). When CP26 content per PSII was at least 50% of that of Wt, the NPQ induction capability was saturated and similar to Wt. These data confirm the crucial role of CP26 in the mechanism underlying NPQ in *C. reinhardtii*, indicating that half of the CP26/PSII content observed in Wt is sufficient to ensure the complete activation of this photoprotective mechanism.

**Figure 8.**
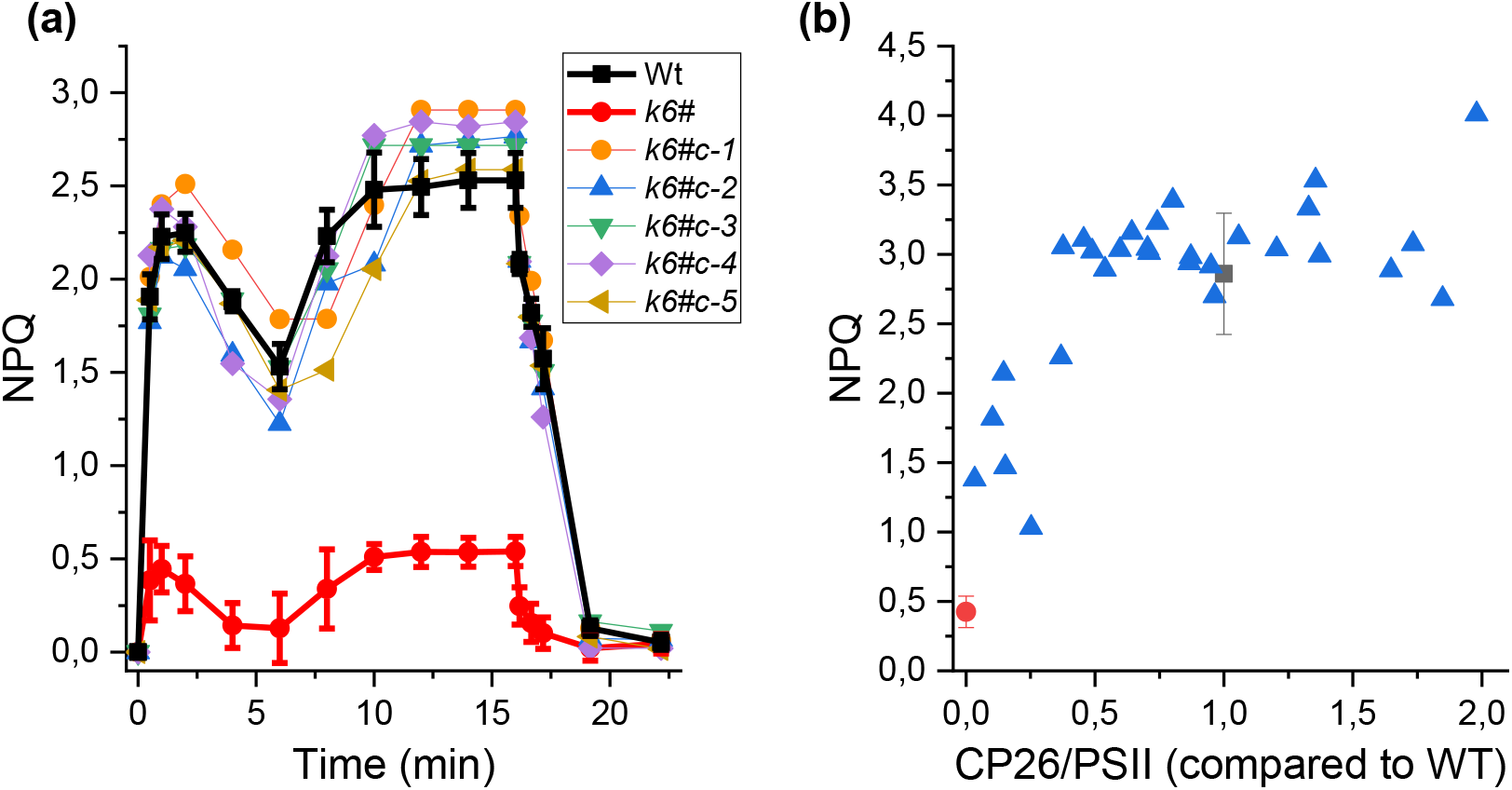
Nonphotochemical quenching in *k6#* complemented line. (a) NPQ using an actinic light of 1,200 μmol m^−2^ s^−1^ in five *k6#* lines complemented with CP26. The selected lines (named *k6#c1-5*) accumulate CP26 to the same level as the wild-type. Wil-type (Wt, black) and *k6#* (red) NPQ traces was added as reference. (b) Correlation between the amount of CP26 and the NPQ reached after 15 minutes of illumination with actinic light at 1200 μmol m^−2^ s^−1^. Different *k6#c* lines are indicate as blue triangles, Wt as a black square and *k6#* as a red circle.

## DISCUSSION

In a previous study, we analyzed the effect of the absence of CP29 or both CP26 and CP29 in *C. reinhardtii* (Cazzaniga *et al*., 2020). The simultaneous absence of CP26 and CP29 affected both photosynthetic efficiency and photoprotection with an almost complete impairment of NPQ induction (Cazzaniga *et al*., 2020). However, a comprehensive study of the specific role of CP26 protein in light harvesting and NPQ mechanism still lacked so far. The CRISPR/Cas9 approach attempted in our previous work to obtain a single *cp26* mutant indeed resulted in the absence of both CP26 and CP29 (Cazzaniga *et al*., 2020), leading to the suggestion of a possible destabilization of CP29 accumulation in the absence of CP26. Here we confuted this hypothesis in two independent ways: i) we successfully complemented the double *k69* mutant with CP29 only and ii) we obtained a single *k6#* mutant by selecting new target regions for Cas9 activity (Supporting Information Table S1). In both cases, CP29 subunit was accumulated to the WT level. On the base of our findings we can thus suggest that the absence of CP29 in the lines generated by CRISPR/Cas9 targeted to *CP26* gene in our previous work (Cazzaniga *et al*., 2020) was likely related to some off-target mutations at the level of *CP29* gene or to some other genes which control the expression of CP29 at translational or post-translational level. These mutations are, however, extremely difficult to be detected and properly evaluated. Here, the availability of a single *k6#* mutant and its complemented lines allow to investigate the effects of the lack of this monomeric antenna on photosynthesis and cell physiology.

In the absence of CP26, Fv/Fm was lower, due to an increase of the F_0_ fluorescence, and the low temperature fluorescence showed a shoulder around 680 nm that represent defectively connected LHCII (Fig. 3). Similar results were obtained in the absence of CP29 in *k9* mutant and were even more evident in the double mutant *k69* (Supporting Information Fig. S5a). These data imply a reorganization of PSII supercomplex in the absence of monomeric antenna subunits, with a reduced efficiency of energy transfer from a portion of the antenna system to the core. Accordingly, time-resolved fluorescence analysis on whole cells demonstrated a reduced energy transfer from PSII antenna complexes to the RC in the absence of CP26, CP29 or both (Fig. 3c, Supporting Information Fig. S5b). Moreover, in *k6#* the LHCII content was strongly increased compared to Wt (Fig. 2) similarly to previous result on the single *k9* mutant, even if a further increased LHCII content was observed in the *k69* double mutant strains (Cazzaniga *et al*., 2020). It is important to note that CP29 subunit stoichiometry per PSII was not changed in *k6*, while *k9* mutant partially compensate the absence of CP29 by a 50% increase of CP26 (Supporting Information Fig. S3). Detailed electron microscopy analysis would be required to assess if the extra CP26 produced in *k9* mutant partially substitute CP29 in PSII supercomplexes. These results indicate the possibility of a compensatory mechanism that increases the amount of antenna to counteract a less efficient excitation energy transfer to PSII RC. This compensatory increase in LHCII was not entirely successful for all mutant strains, as can be seen by appearance of fluorescence signatures at 77K related to “unconnected” LHCII in the absence of CP26, CP29 or both. Moreover, in *k6#* the functional antenna size was similar to the Wt even if the amount of LHCII was almost doubled (Fig. 3). The reduced energy transfer efficiency in the absence of CP26 influenced the photosynthetic activity of PSII, causing reduced light dependent oxygen evolution, especially at low-medium irradiances (Fig. 4, Supporting Information Table S3). However, the similar Pmax of Wt and *k6#* mutant suggests that proper PSII photochemical activity can be restored even in the absence of CP26, providing increased photons to counterbalance the reduced light harvesting efficiency observed in the mutant. These results differ from that of the mutant strains without CP29 (*k9* and *k69*), where the oxygen evolution rates were always lower than Wt, resulting in reduced Pmax. The altered photosynthetic activity in absence of CP26 also affected biomass accumulation when grown in limiting light condition, where the *k6#* exhibited a lower cellular density and the growth rate was reduced compared to Wt. However, a more severe reduction in growth rate could be observed in the case of *k9* and *k69* (Supporting Information Fig. S6), consistent with the lower photosynthetic activity measured in the absence of CP29 or both CP26 and CP29 compared to the case where only CP26 was missing. When mutant strains were grown at higher light intensities (700-800 μmol m^−2^ s^−1^) similar growth curves were observed for Wt and mutant strains, suggesting that in these conditions light harvesting and PSII activity was not limiting the growth. All these data point out that both monomeric antenna proteins are important for efficient light harvesting in *C. reinhardtii* but CP29 have a prominent role. These functional differences correlate with the position of the two antennae in the PSII supercomplex: CP29 is located in the inner layer of the antenna and interact with trimers S-LHCII, M-LHCII and marginally with additional N-LHCII while CP26 is in a more external position and interact only with S-LHCII that is still connected to the RC from the other side through CP29 (Shen *et al*., 2019). Due to this organization, the absence of CP29 had a stronger impact on PSII antenna complex and on efficiency of light energy direction toward the core complex.

Photosynthetic organisms evolved specific strategies to maximize light harvesting in low light condition and avoid excess energy absorption in high light. In the case of the model organism for green algae, *C. reinhardtii*, NPQ induction requires the presence of LHCSR proteins which upon thylakoid lumen acidification, are activated through their pH sensing residues (Liguori *et al*., 2013; Ballottari *et al*., 2016). LHCSR proteins need to interact with other antenna proteins to efficiently redirect and quench the excess light energy absorbed. (Liguori *et al*., 2013; Ballottari *et al*., 2016). Alternatively, it cannot be excluded that LHCSR can trigger a conformational change in other antenna subunits, leading to the formation of additional energy dissipation sites (Tokutsu & Minagawa, 2013).

Previous works demonstrated by electron microscopy the interaction of LHCSR3 dimers with CP26 and LHCII subunits but the specific functions of the different antenna proteins in the NPQ process has not been fully elucidated yet. In the case of LHCII, the absence of LHCBM1 (Elrad *et al*., 2002) or downregulation of LHCBM4-6-8 subunits (Girolomoni *et al*., 2017) caused a reduction of NPQ induction, suggesting these subunits as potential partners for LHCSR3. Our previous findings demonstrated that in the absence of monomeric CP26 and CP29 subunits, NPQ induction was essentially impaired in *C. reinhardtii* (Cazzaniga *et al*., 2020). Here we can narrow down our knowledge, demonstrating a specific function for CP26 in NPQ. The NPQ was strongly reduced in *k6#* mutant at all the light intensities tested, even with strong actinic light (Fig. 6, Supporting Information Fig. S8). The absence of CP29 in *k9* also caused a decrease in NPQ compared to Wt but to a lesser extent compared to that in *k6#* while a more dramatic NPQ phenotype was observed in the absence of both CP26 and CP29 subunits in *k69* mutant (Supporting Information Fig. S8). The impaired NPQ mechanism in *k6#* could be due to a decreased LHCSR/LHCII ratio which was reduced in *k6#* compared to that in Wt. The relatively reduced LHCSR/LHCII stoichiometry in *k6#* could indicate a lower possibility to quench the increased amount of trimer in the absence of CP26 resulting in a lower level of NPQ. However, in the *k9* the LHCSR/LHCII ratio was similar to that in *k6#* (Cazzaniga *et al*., 2020) but the NPQ induction was less affected. Thus, reduced NPQ in the absence of CP26 may not be due to the alteration of LHCSR/LHCII ratio. Different lumen acidification could also be excluded as possible explanation for reduced NPQ observed in *k6*, according to the similar NPQ impairment observed upon induction of NPQ in the dark by artificial lumen acidification. The specific role of CP26 in NPQ was confirmed by the genetic complementation of *k6#* mutant (Fig. 8). Interestingly, accumulation of CP26 per PSII to a 50% level compared to the Wt case was sufficient to fully restore NPQ. PSII supercomplexes are included in the tightly stacked grana of the thylakoid membranes and single PSII can pair up with near complexes or connect with complexes across the stromal gap (Albanese *et al*., 2017). It is possible to speculate that these close interactions may allow CP26 to interact with different complexes, reducing the amount needed of this subunit to transfer excitation energy from PSII to the quenchers, as LHCSR proteins. These data confirm the fundamental importance of monomeric antenna subunits in the quenching mechanism (Cazzaniga *et al*., 2020) but indicate CP26 as the monomeric antenna protein with a prevalent role in NPQ. These results are consistent with previous evidence that points out to CP26 as docking sites for LHCSR3 on PSII supercomplexes (Kim *et al*., 2017; Semchonok *et al*., 2017). LHCSR proteins are not constitutively expressed in *C. reinhardtii* but only upon high light exposure stress (Tokutsu & Minagawa, 2013): being in an external position, CP26 could represent a more accessible point for a direct interaction with LHCSR3, while CP29 is clamped between the core and S- and M-LHCII (Drop *et al*., 2014; Cao *et al*., 2020). Monomeric CP26 could thus be essential for the energy transfer from the antenna complex to LHCSR subunit. The dramatic difference in NPQ between *k6#* and *k9* could be influenced also by the change of expression level of the remaining monomeric antenna in the single mutants. While in Wt and *k6#* the relative amount of CP29 is similar, *k9* shows an over-accumulation of CP26 protein, with an increase of ∼ 50 % with respect to Wt. This increase in CP26 could partially compensate for the absence of CP29. This implies that CP26 could interact also with PSII core in the region where normally CP29 resides improving LHCII connection. This compensatory mechanism has never been observed in *Arabidopsis thaliana*, where CP29 binding site cannot be occupied by other subunits (de Bianchi *et al*., 2011). In *A. thaliana*, where the absence of CP26 had no effect on the quenching mechanism, the simultaneous deletion of all monomeric subunits reduced the light harvesting capacity but had a minor role in NPQ (de Bianchi *et al*., 2008; Dall’Osto *et al*., 2020). The data from this study further confirm the differences between LHCSR-dependent NPQ observed in *C. reinhardtii* and the PSBS-dependent NPQ in vascular plants. In both cases, LHCSR proteins and PSBS need to interact with LHC proteins to trigger NPQ (Niyogi & Truong, 2013; Sacharz *et al*., 2017): a possible different affinity of LHCSR and PSBS for the specific LHC antenna could explain the different phenotype observed in LHC mutant in *C. reinhardtii* compared to *A. thaliana*.

In conclusion, these results point out a distinct role of the two monomeric antenna subunits in *C. reinhardtii*. While CP29 is fundamental for efficient light harvesting and PSII activity, CP26 plays a major role in NPQ induction. It is important to note that a small NPQ induction at very high light could still be observed in the absence of both CP26 and CP29: this residual NPQ was probably due to the interaction of LHCSR1 or LHCSR3 with LHCII subunits, as previously reported (Kim *et al*., 2017; Semchonok *et al*., 2017). These data could be useful to further understand the different NPQ molecular mechanisms in microalgae compared to vascular plants and to identify targets to tune this mechanism to increase productivity under different light regimes.

## ACKNOWLEDGEMENTS

The research was supported by the ERC (European Research Council) Starting Grant SOLENALGAE (679814) to M.B. and by Basic Science Research Program (NRF2020R1A2C2011998) and Carbon to X Project (2020M3H7A1098294) of the National Research Foundation (NRF) of Korea, funded by the Korean government. to E.J.

## AUTHOR CONTRIBUTIONS

MB designed the research. EJ coordinated the research activity for generation of mutant strains and contributed to experiment design. CD coordinated the research activity for time resolved fluorescence analysis of whole cells. SC, MK, SS, FP and MP performed experiments. All the authors analyzed data. SC, EJ and MB wrote the paper. All the authors discussed the results, contributed to data interpretation, commented on the manuscript and approved its final version.

## Authors declare no conflict of interest

## SUPPORTING INFORMATION

**Methods S1. CRISPR-Cas9 genome editing and acid induced quenching Figure S1. Expression of CP29 in CP26/CP29 double mutant background**

**Figure S2. Screening and verification of CP26 knock out mutants**.

**Figure S3. CP26 immunotitration in *k9* mutant**.

**Figure S4. F**_**0**_ **and F**_**m**_ **fluorescence normalized to chlorophyll content**.

**Figure S5. Photosynthetic parameters in *k9* and *k69-***

**Figure S6. Growth curves for *k9* and *k69***.

**Figure S7. Immunotitration of LHCSR proteins per PSII**.

**Figure S8. Nonphotochemical quenching at different actinic lights**.

**Figure S9. Nonphotochemical quenching in *k9* and *k69* mutants**

**Figure S10. Acid induced quenching in 4a+ and *npq4 lhcsr1***

**Figure S11. Acid induced quenching in *k9* and *k69***.

**Figure S12. *k6#* mutant complementation**.

**Figure S13. *k6#* line complemented with CP26**.

**Table S1. Sanger sequencing of the CP26 knock-out mutants**.

**Table S2. Fluorescence lifetimes of whole *Chlamydomonas reinhardtii* cells**.

**Table S3 Light curve parameters and respiration rates**.

