## Supplementary material for "Photosystem II monomeric antenna CP26 has a key role in Non-Photochemical Quenching in *Chlamydomonas reinhardtii*": Fig. S

**Methods S1. CRISPR-Cas9 genome editing and acid induced quenching**

*CRISPR-Cas9 genome editing*

The sgRNA sequence for the CP26 mutant generation using CRISPR-Cas9 was designed by Cas-Designer (http://www.rgenome.net/cas-designer) and selected as 5′- CCCGAGGGCCTGCAGGCCAACGG -3′ (Figure 1a). To increase the efficiency and fast selection of CP26 mutants, insertion and expression of hygromycin-resistance (HygR; aphVII) gene cassette was combined. 100 µg of purified Cas9 protein (Cas9 expression plasmid: pET-NLS-Cas9-6xHis (Plasmid #62934)) and 70 µg of sgRNA synthesized by using GeneArt™ Precision gRNA Synthesis Kit (ThermoFisher, MA, USA), were mixed to form the RNP complex and co-transformed with aphVII gene expression cassette. After transformation, cells were plated on TAP medium containing 1.5% agar and hygromycin (25 µg/ml). To screen CP26 knock-out mutants, the colonies grown onto the selection plate were confirmed by genomic PCR using the target-specific primer set (forward: 5′-CTTTCAGTCAGCTTGCATGG -3′, reverse: 5′-AGGGGGTTGGACACAATG -3′). The PCR products were sequenced using Sanger sequencing by Macrogen Inc. (Seoul, South Korea).

*Acid induced quenching*

Chlamydomonas reinhardtii cells were first adapted to high light for two days to induce LHCSR. Adapted cells cells were centrifuged (3 min, 1000 g), resuspended in water at the density of $10^6$ cells/ml and were acidified by adding 5 mM acetic acid. Fluorescence was measured with a PAM-101 fluorimeter (Walz, Effeltrich, Germany): maximum fluorescence emissions were measured applying saturating pulses, before ($F_m$) and after ($F_m^a$) decreasing the pH with acid solution. NPQ was calculated upon reaching steady state fluorescence levels as $(Fm-Fm^a)/Fm^a$

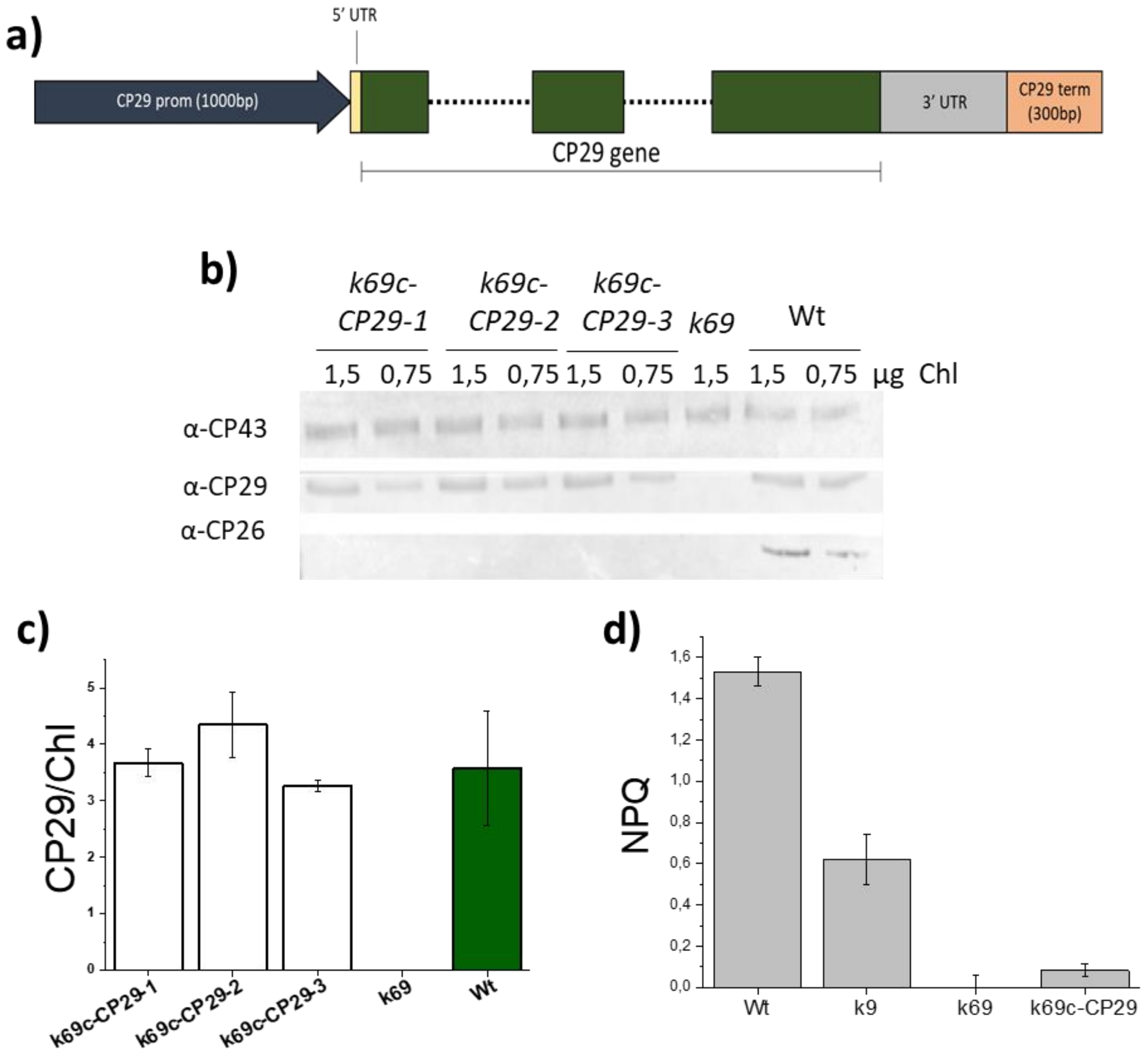


**Figure S1. Expression of CP29 in CP26/CP29 double mutant background.** (a) genomic sequence scheme of the CP29 (Cre17.g720250) gene according to *Chlamydomonas reinhardti*i v 5.6 genome. 5' and 3' UTR are colored in yellow and gray respectively, while CP29 coding sequence is in green with introns represented as dotted lines. The putative promoter (1000 bp upstream) and terminator (300 bp downstream) are reported in blue and orange respectively. (b) Immunoblot analysis of CP29 and CP26 expression in Wt, double mutant *k69* (deprived of both CP26 and CP29) and *k69* complemented with CP29 gene (*k69c-CP29*). Chlorophyll (Chl) loading is reported. CP43 was also analyzed on the same filter as loading control. (c) Densitometric analysis of western blot results reporting the CP29 content in the strains analyzed in b). d) Non-photochemical quenching (NPQ) of Wt, single mutant on CP29 (*k9*), double mutant *k69* and CP29 complemented lines in *k69* background (*k69c-CP29*). The NPQ values were measured on dark adapted cells after 90 seconds of illumination with 1200 µmol $m^{-2}s^{-1}$ actinic light. Differently from the other NPQ measurements reported in this work, here no far-red adaptation was applied before exposure to actinic light. Error bars are reported as standard deviation (n=3).

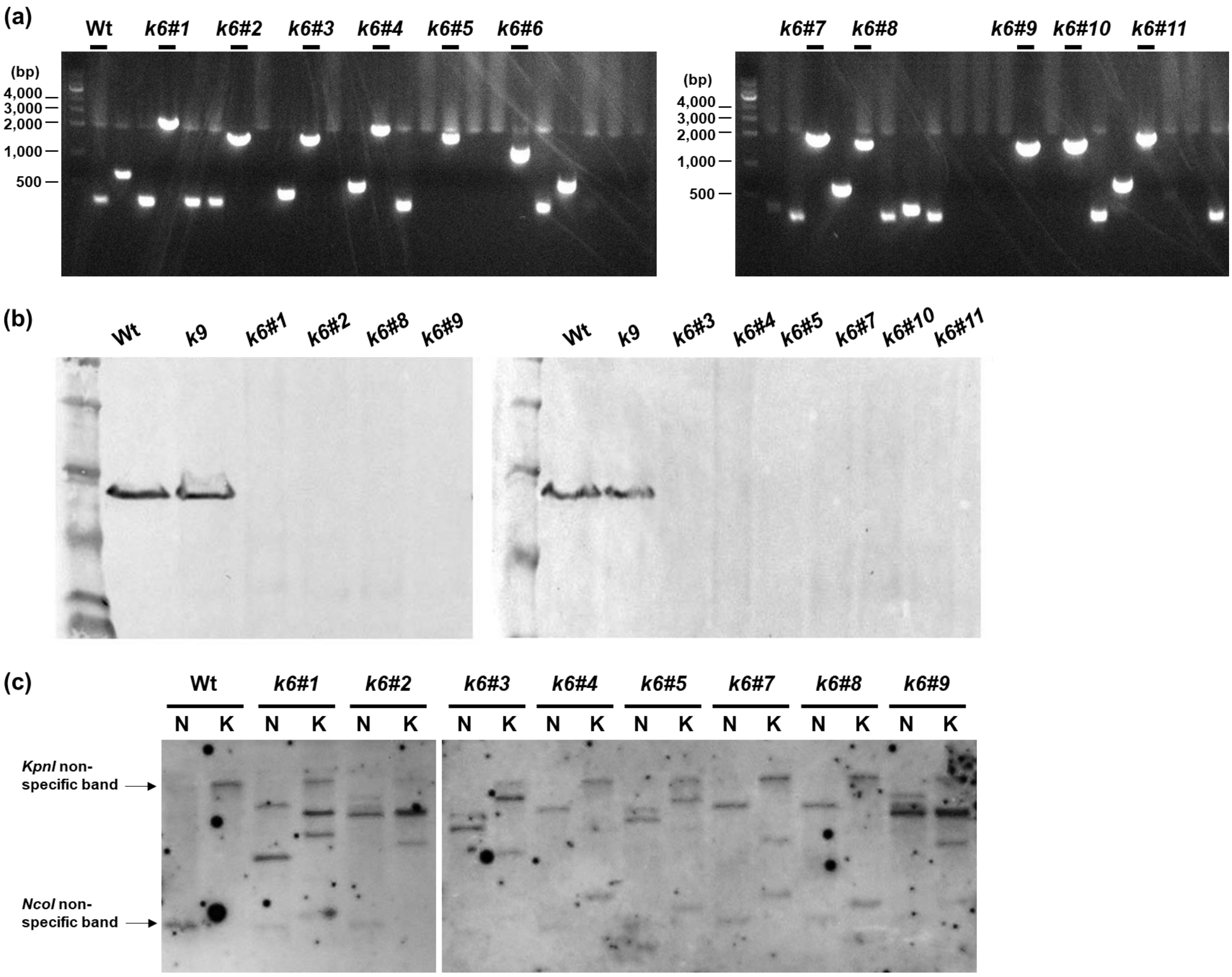


**Figure S2. Screening and verification of CP26 knock out mutants.** (a) Colony PCR using the specific primers adjacent to sgRNA target sites of CP26 gene in wild type and putative CP26 knockout mutants (*k6#*). (b) Immunoblot analysis of selected knock out mutants of CP26. Total protein extracts of *k6#* lines were loaded on SDS-PAGE gel and checked for the presence of CP26 with a specific antibody against CP26. Total protein extracts from Wt and *k9* line (mutant without CP29) were also added to the external lanes as loading controls. (c) Southern blot on selected CP26 knock-out (*k6#*) mutants. Extracted genomic DNA was digested by *Kpn*I and *Nco*I and blotted on a nylon membrane. The *aphVII* probe was used for detection of DNA integration.

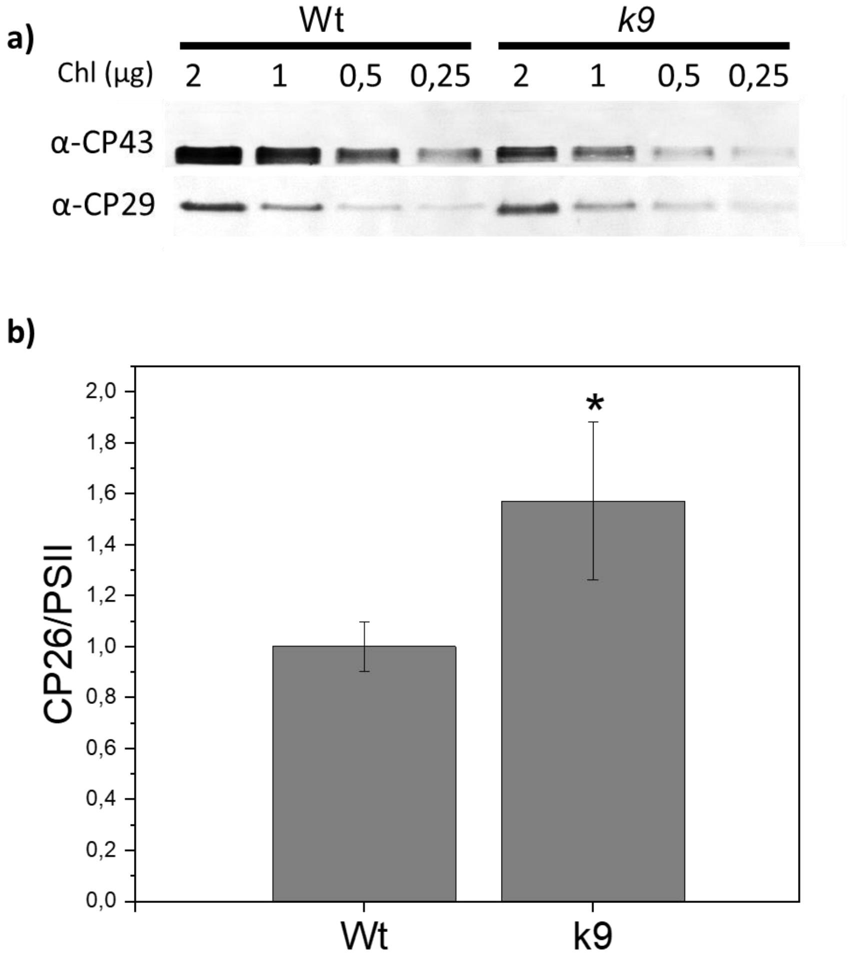


**Figure S2. CP26 immunotitration in *k9* mutant**. Immunoblot analysis of CP26 content in the *k9* mutant (a). Densitometric analysis of the immunoblotting results were used to determine the CP26/PSII ratio by using CP43 as a proxy for Photosystem II (PSII). Error bars are reported as standard deviation (n=4). Values that are significantly different in *k9* mutant compared to Wt are marked with * (Student's t-test, P < 0.05).

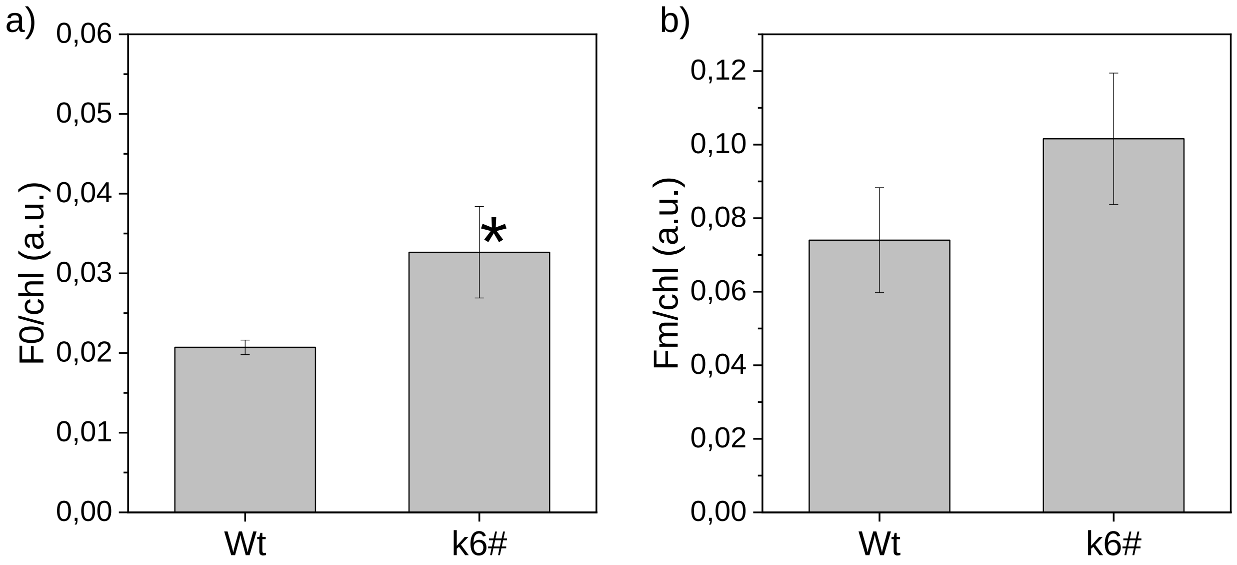


**Figure S3. $F_0$ and $F_m$ fluorescence normalized to chlorophyll content.** Dark-adapted wild type (Wt) and *k6#* cells were excited with the same PAM light setting and minimal ($F_0$) and maximal ($F_m$) were recorded. After the measure Chl were extracted and quantified from all the samples. $F_0$ (a) and $F_m$ (b) were normalized Chl content). Error bars are reported as standard deviation (n=6). Values that are significantly different in *k6#* mutant compared to Wt are marked with * (Student's t-test, P < 0.05).

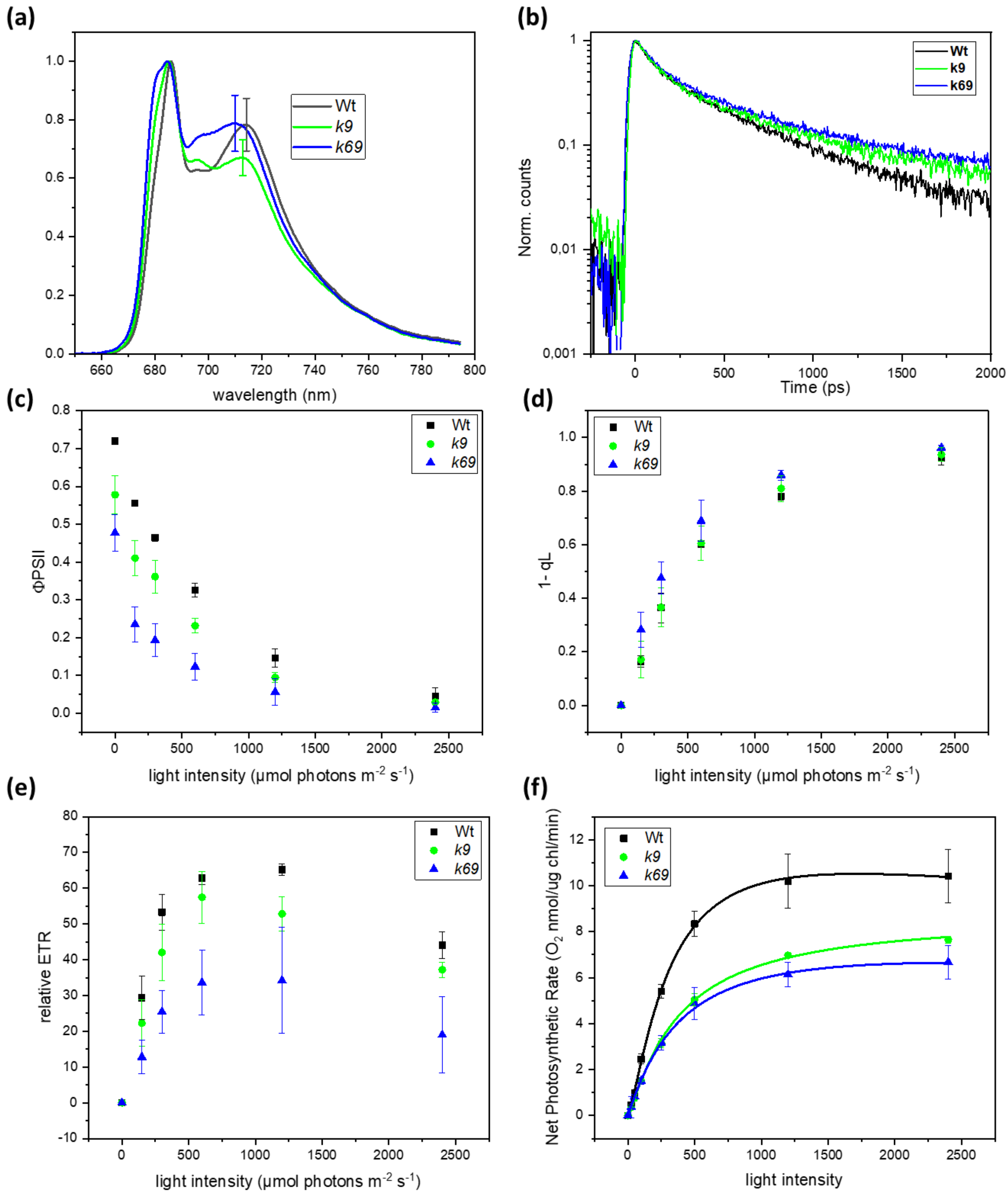


**Figure S4. Photosynthetic parameters in *k9* and *k69*.** (a) Low-temperature fluorescence emission spectra, (b) Time-resolved fluorescence decays, (c) PSII operating efficiency (ΦPSII), (d) relative electron transport rate (ETR), (e) 1-qL (reduced $Q_A$ fraction), and (f) photosynthetic O2 evolution for wild type (Wt black), *k9* (green) and *k69* (blue). Error bars are reported as standard deviation (n=4).

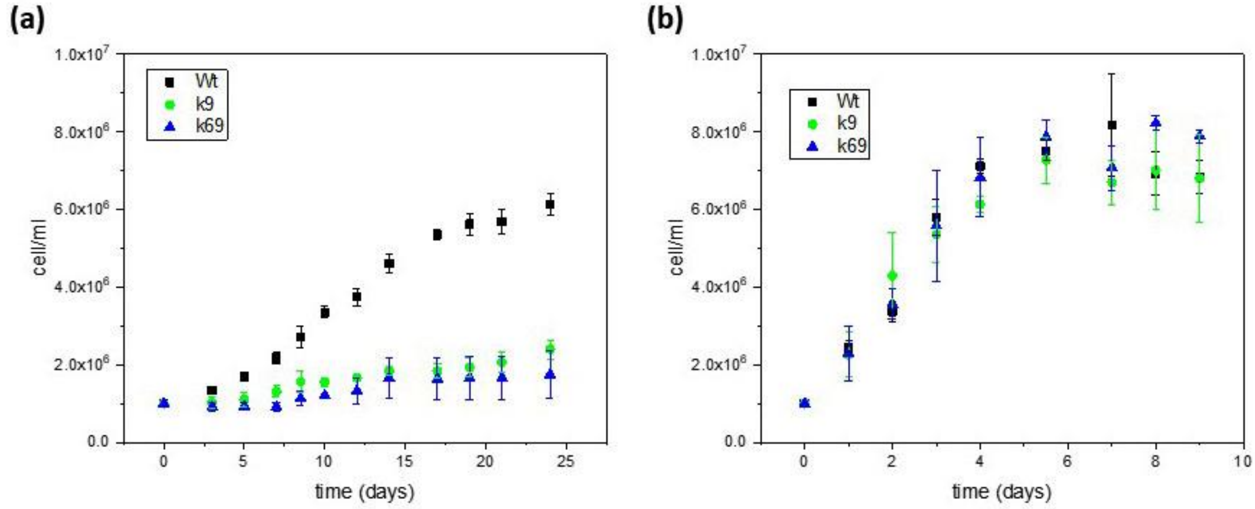


**Figure S5. Growth curves for *k9* and *k69*.** Growth curves of wild type, *k9* and *k69* at (a) low (30-40 µmol $m^{-2}$ $s^{-1}$, LL) and (b) high light (1200 µmol $m^{-2}$ $s^{-1}$, HL). Growth curves were fitted with Hill equation. Error bars are reported as standard deviation (n=3).

a)

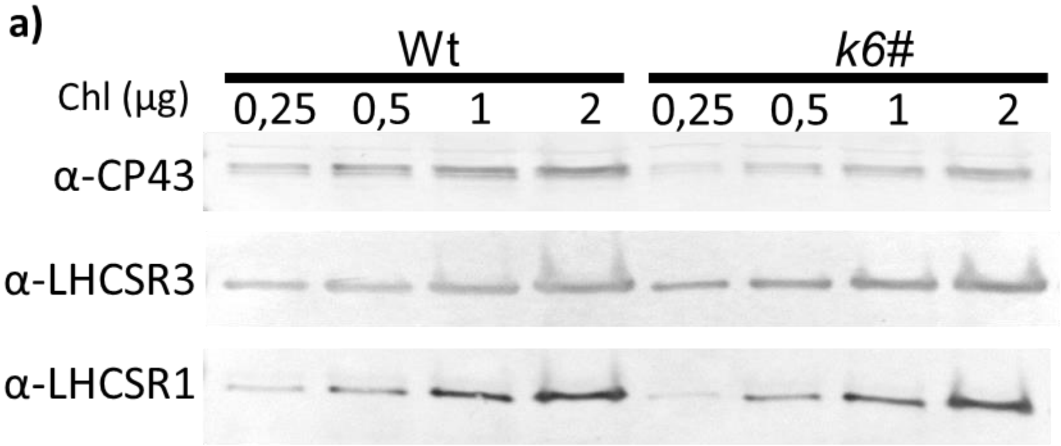


b)

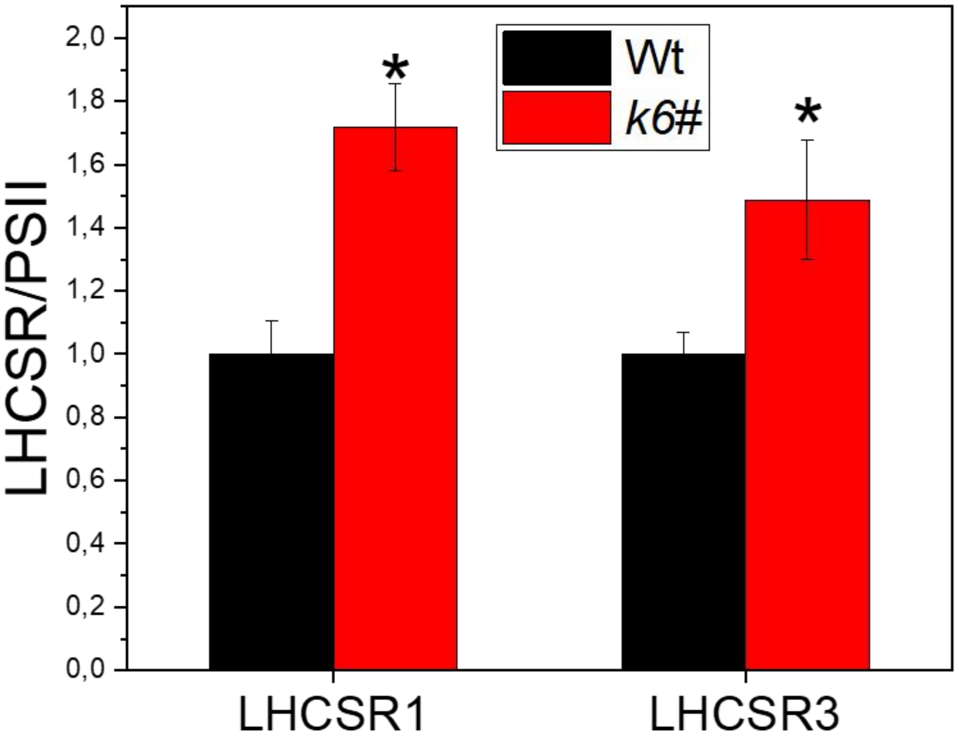


**Figure S6. Immunotitration of LHCSR proteins per PSII.** (a) Western blot for immunotitration analysis on high light adapted cells of wild type (Wt) and *k6#* with specific antibodies against LHCSR3 and LHCSR1. 0.25 0.5, 1 and 2 µg of Chl were loaded on the different lanes as indicated. (b) LHCSR1 and LHCSR3 quantifications by immunotitration analysis. LHCSR/CP43 ratios were normalized to the Wt value. Error bars are reported as standard deviation (n=3). Values that are significantly different in *k6#* mutants compared to Wt are marked with * (Student's t-test, $P < 0.05$).

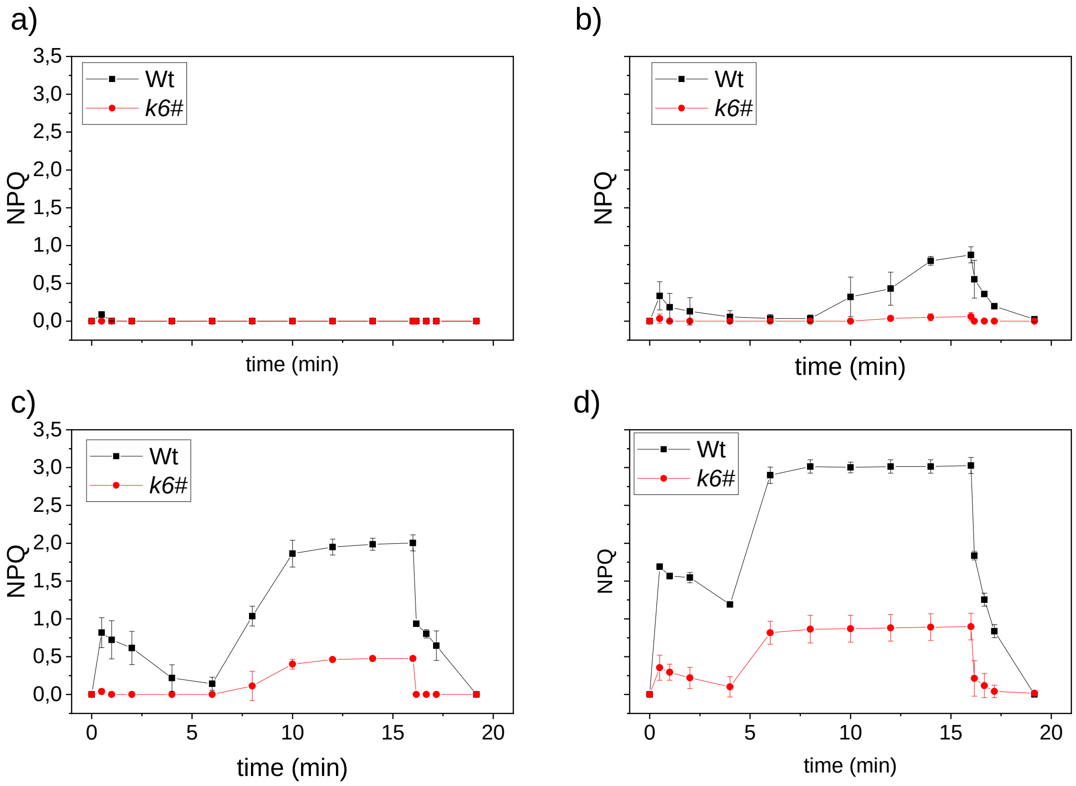


**Figure S7. Nonphotochemical quenching at different actinic lights.** NPQ kinetics of wild type (Wt, black) and *k6#* (red) cells using actinic lights of (a) 150, (b) 300, (c) 600, and (d) 2400 µmol photons $m^{-2}$ $s^{-1}$. Error bars are reported as standard deviation (n=4).

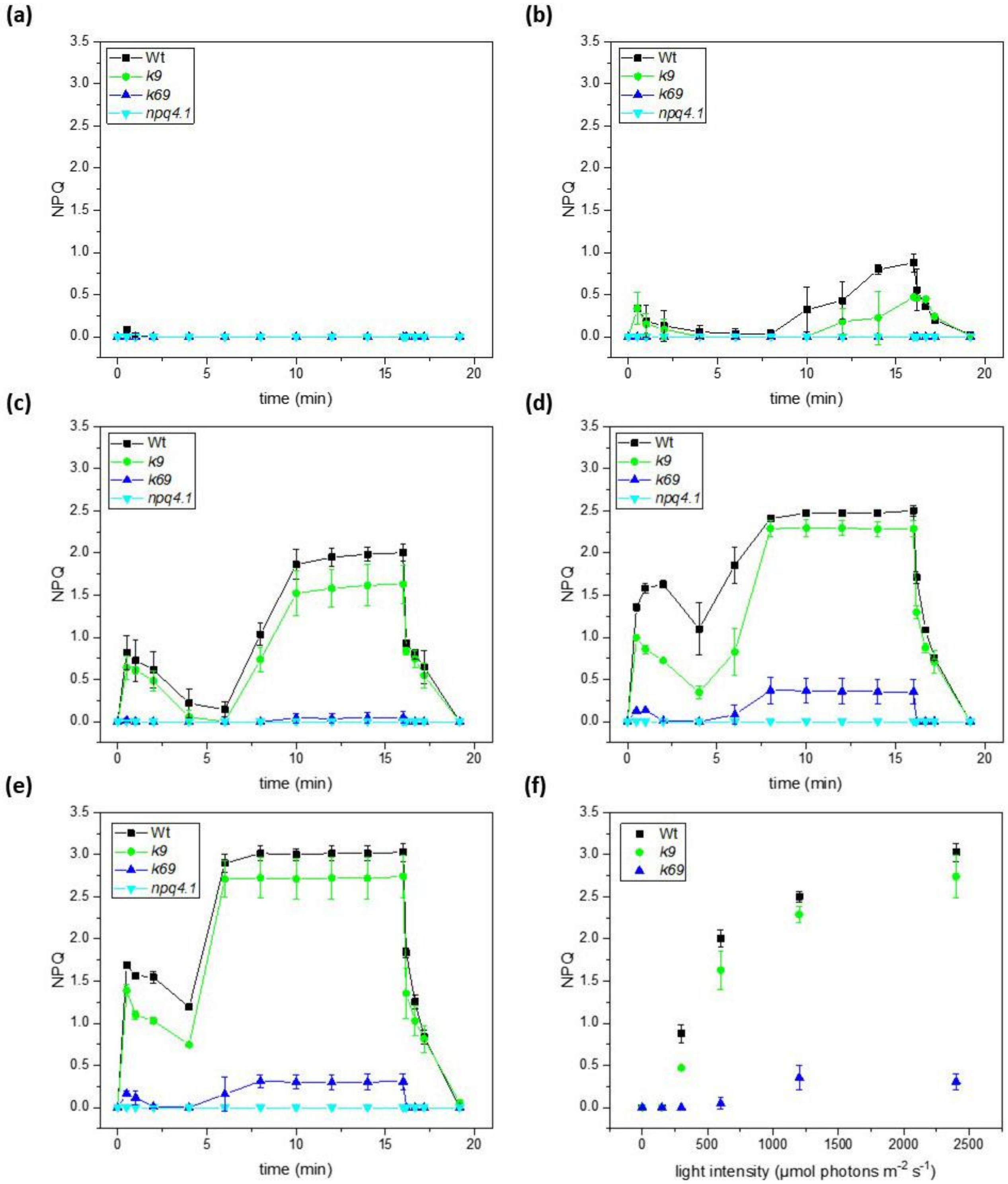


**Figure S8. Nonphotochemical quenching in *k9* and *k69* mutants.** Measurement of NPQ kinetics on wild type (Wt, black), *k9* (green), *k69* (blue) and *npq4 lhcsr1* (npq4.1 cyan) cells using actinic lights of 150 µmol (a), 300 (b), 600 (c), 1200 (d) and 2400 photons $m^{-2}$ $s^{-1}$ (e). (f) NPQ value after 15 min of illumination at different actinic light intensities. Error bars are reported as standard deviation (n=4).

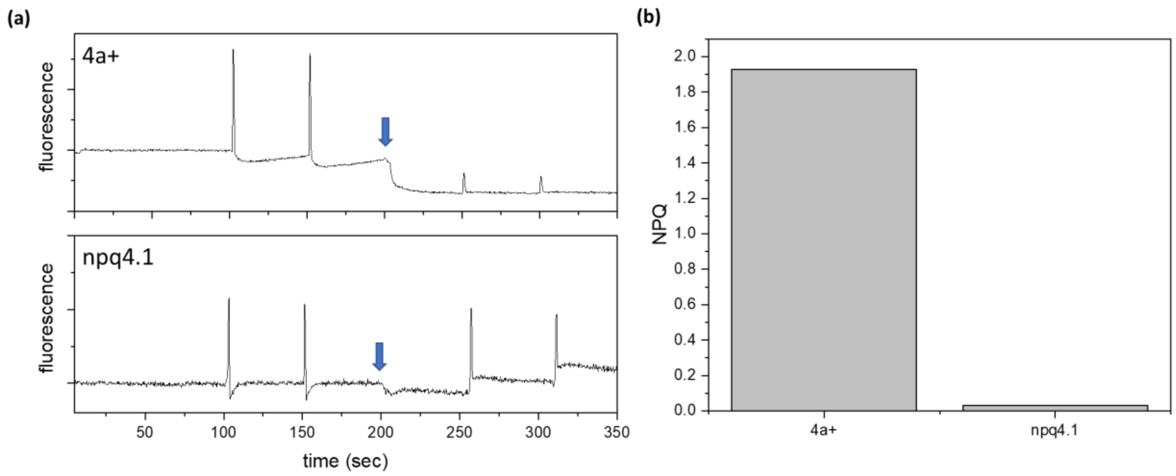


**Figure S9. Acid induced quenching in 4a+ and *npq4 lhcsr1*.** (a) PAM Chl fluorometer trace of 4a+ (above) and *npq4 lhcsr1* (herein named *npq4.1*, below) before and after acidification with acetic acid. The blue arrow indicates when the acid was added. Fluorescence was recorded in the dark with two saturating pulses before and two after the acidification. (b) NPQ is calculated for each trace as (Fm-Fm$^a$)/Fm$^a$ where Fm and Fm$^a$ are the fluorescence before and after acidification.

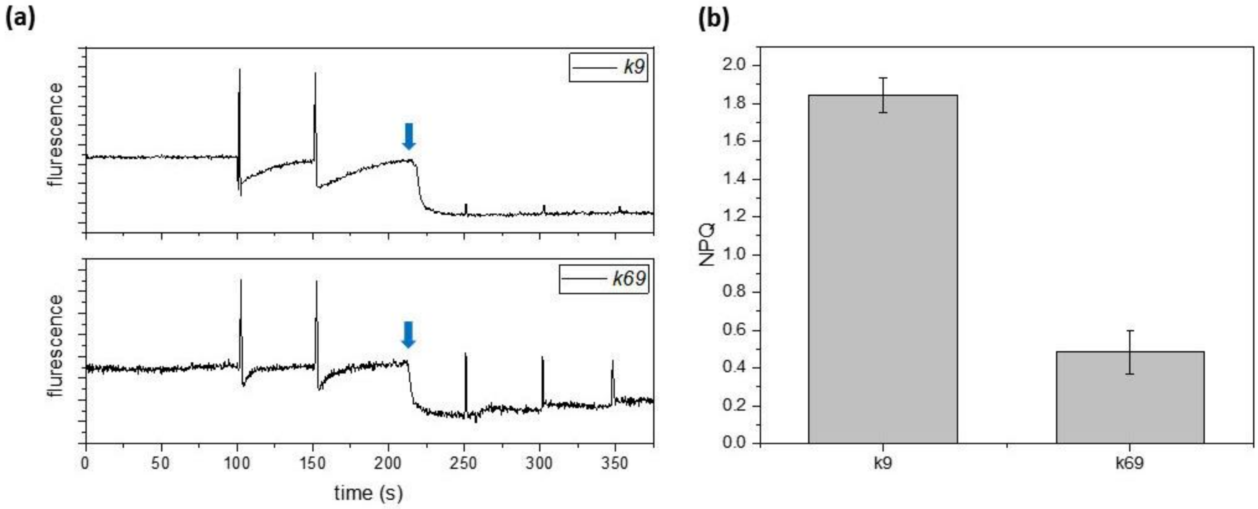


**Figure S10. Acid induced quenching in *k9* and *k69*.** (a) PAM Chl fluorometer trace of *k9* (Wt) (above) and *k69* (below) before and after acidification with acetic acid. The blue arrow indicates when the acid was added. Fluorescence was recorded in the dark with two saturating pulses before and two after the acidification. (b) NPQ is calculated for each trace as (Fm-Fm$^a$)/Fm$^a$ where Fm and Fm$^a$ are the fluorescence before and after acidification.

(a)

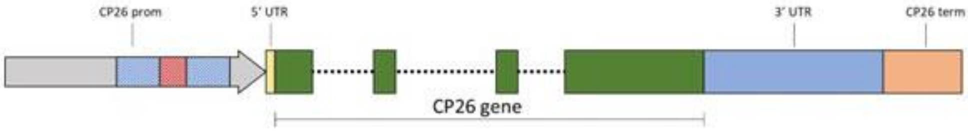


(b)

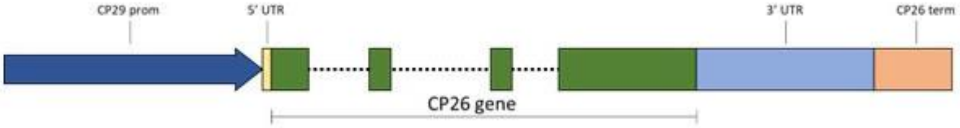


(c)

CGTCCACATTTACCGGAGCGTGCCGGCGAAGTGCCGGAGCTCGAAGGGCCAACGCACCCTAACGCACTTTCGGCGGCC
TTTCGAGTGCCCCTCTGTACCCTGCTGCCTGCCCATAAGCATGGTCGCAGATGAAAGACGGGCAAGACACGATTATCC
TGCAGGCAATTGCCGGCGCGAGCTTGGGGCGCCCCTTCAGCGTCCCATCGGCGGTCGCTTTTTGCCCCGGTGTCGCCG
TTCCTGGTTCTCGGCAGCCCAAGATAATTTAATCTAGTAGTAATAATCATGTGCAGCGTTGTGGCAGCTGCCCCCAAA
GGAAACTGTGGCGGGAAGCGCCCCAGTCGCGCAAGCTTATCGCTCGGTCGCGCGTCGGGGCCACCCTGAAGACCCTGA
ATTATTTGTGCGACAATATAGCAGCCACTTCTTTTCATTTGAATGGTTTTCCGACAGCGAGAGGCGCGAAATATTATG
GTTGCGGGGGTTGCGGGGAGGGGGAGGGGGAGGGGGAGGGGGAGGGGGAGGGGGGAGGGGGAGGGGGGAGGGGGGAGG
GGGAGGGGGAGGGGGAGGGGGANNNNNNNNNNNNNNNNNNNNNNNNNNNNNNNNNNNNNNNNNNNNNNNNNNNNNNNN
NNNNNNNNNNNNNNNNNNNNNNNNNNNNNNNNNNNNNNNNNNNNGGGAGGGGGAGGGGGAGGGGGAGGGGAGGGGAGG
GGAGGGGAGGGGAGGGGAGGGGAGGGGAGGGGCGGGGCGGGGCGGGGCGGGGCGGGGCGGGGCGGGGAGGGGAGGGGC
GGGGCGGGGCGGGGCGGGGCGGGGAGGGGAGGGGCGGGGCGGGGCGGACAAATAGGTCAGCAAATGGATGAACATGAC
CGCAAATTGATAATCATACCTGGCTTGCAAGCTCGCGCCCAGCGAGATGGAGTACGGACGATGGAGATCTGGCCGCGA
TTGGCGAGCCGGGCAAGAAAAACAGCCGAGCGCTGCATATAACACTTGTCACACCGTCGACCTT

**(d)**

TCGCATACACTCGGCGCCGCGCAACTGTTGCATCCCCAACCTGCCCCATCACACCCAACACAGCCCGCCAACACTCTCCTCCCGCCACCACCATCAGCAC
CTGCCCGCCTGGATGCACAGGCCTCAAGGTCAGAACCGGTGCTTGCCCGCTCGTGCGTGCTATCGGTGGACTCGACCCTATCATTGGGCTCGGGCAAGTA
TCTCCATTCCCCGTGCCCCTCCGTGCCCCGCCAGGCGAGTCGGCCACCCCCCGGATAGCACCAGCCCATGCCCCCAGGCCATAGCTGCCCTGGACTGGTT
CTCACGAATCACCGCTCACGCATGTTAAACAGCCAAGCAGTCACATTGCCAACACGAGCGCGTGTGCACCCGAGAACCAGAATACAGCGATGCACTTGCG
CACCAGCTCAAGCCAAAAAGATGGCGGTTCCAGTTTTGCGCACTGCACCATCGTCCCGAGTAGGCCGCCTTCCTTGACACTGAGAGCGAGACAGCGAGAC
TTGGCGCCGATGGGTGTTGCGGTGTTGCACGTTTGCTTGCTGCGCACACGCTTGCCTGCCCGGCACCCCTGCTTAAGTCGGTATGCACCTATAAGTGTCG
AAAACTGAACGCCATATTGAGGCGACGATACGGAGCCAGCGCGCAAGCGCATGTCGGGCCATGTAGAACCTACACTTGCTCGCTTCCCCGTCGCTACCTA
ACGCGAAGCAATGCATTAACATGCGCCTAAGTGTTGAAGTTATTGTAACAAGCACGGGTCCATTTTGCGCGCCAAAAGGGCCCCAGCTCGCTGCAAGGGG
CGCTCGCGGCACGGTCGCTGGTGGCGCGGGCCAGCGCTCTAGTGAATACAATTCATTTGGCGTCTTGTCGCGGCGCTAGAGGGTCGTTCCGGCATGAGAG
CGCGAAGGAATGAGGCAGATCTGTGTACGGCGGCCCCGAGAAGATGGGGCTTTAGAAAAGGCTCGTCGAGCCACCGTTTCGTTAGTCTTTGTAGCACATA
GTTCGTTCAGTCACTTGAACAGCAACACCATGCAGATCCAGGCTCTGTTCAAGAAGACCGGCGCTAGCGCCCCCGCCAAGAAGGGCACTGCCTCCACCAA
GGTGGTCAAGCCCTCCAAGGCCGGCGGCAAGGCCACCCGCGGCTGGCTGGGCGGCCAGGGCGGCGCTGCCGACCTGGACAAGTGGTACG~~GTGAGTCGCCG~~
~~GGCGCGATCCTGCTGGGTTGAGCGACCGCATGGCGAATGTGCTAGGGCCATCCCTTGCTTCCCTTGTGCAAGTGGCCCGCCAAACCGCTGGTTGGCTCG~~
~~TAAATGGTGTTGTAAAGCTTCTTCGGGCCTGATCGAATGGCAAACGTCGCCGTGTCGCTGTCGCGCCCCTTCCGCAAGATGCAAGCACCTCACTTATCTG~~
~~GCGTTGCTGATGTTCGCAG~~GCCCCGACCGCAAGCTGTTCCTGCCCAGCGGCCTGTACGACCGCTCGGAGATCCCCGAGTACCTGAACGGCGAGCTGGCTG
GCGA~~GTGAGTCGGGAAATGCGCAGCGCTGGGATACCTGCTCTCGATCAGCGATTTCGTTCACGGAGTGGGGCCTCAAGACTGTTTAGGACCCGTCTCAGC~~
~~CTTGATAGCTTCAGATCAAGGGGTCTGAAGCTGGGTCATTGCGGGCCATCTACGCAGTCGCGATGTGGCACCCACTCGCGGCCTTGGTTCGGCGCTCAAA~~
~~GCATCACTTTGCGAGTCGACTAAAGAATGCCTAGCCAGAACCGTGGGCTGGGACGGCGTCCTTGCGTTACTAGTGATGGCCAGTGCTGCCAGCGTGGC~~
~~TGCCAGCGGACCAGCTATCTCTGTGCCCCGTGCTGCTTTGAGTCGCTTTGTAACCCCTCCCCTCCCCGCTTCACTCTTGTGTCCTTTCTCAG~~CTACGGCT
ATGACCCTCTGGGCCTGGGCAAGGACCCCGAGACCGTGGCCAAGTACCGCGAGAACGAGCTGCTGCACGCCCGCTGGGCCAT~~GTGAGTAGGGGGCTGCTG~~
~~CACTTCTTGTCCTGTGCGCTGGGTTGGTCATGGCGTACAGCTAGGACGGCCATTACAGCTTTCAGTCAGCTTGCATGGCACTCGATTGCGCTGGTGCGCG~~
~~CCGTGCAGCCGCCGTCTGACGTCCCATCCCTTGCCTGCGTCCTCCTGCCCCGCAG~~GCTTGCCGCTGCCGGTATCCTGATCCCCGAGGGCCTGCAGGCCAA
CGGTGCCAACATCAAGGGTGGCACCTGGTTCGAGACCGGCGCTGAGATGCTCAACGGCGGTACCCTGAACTACTTCGCCGTGCCTTGGGGCATTGTGTCC
AACCCCCTGCCCCTGTTCGCCGTCATCGCCGTTGAGGTCGGCCTGATGGGCGCCGTGGAGTTCTACCGCCGCAACGGCACCGGCCCCGCCGGCTACTCCC
CGGGCATTGGCAAGTTCGACTCGTCGGTCTTCGACGGCCTGGACCCCCTGTACCCCGGCGGCCCCTTCGACCCCCTGGGCCTGGCTGACGACCCCGAGGT
CCTGCAGGAGCTGAAGGTCAAGGAGATCAAGAACGGCCGCCTGGCCATGGTCTCCGTGCTGGGCTTCGCCGTCCAGTCCTACGTGACCGGCGAGGGCCCC
TACGCCAACTGGACCAAGCACGTGGCCGACCCCTTCGGCTACAACCTGCTGACCGTCCTGGGCGCCGAGGAGCGCACCCCCACCCTGTAAATGCCTTTTC
CAGCTACAGCCTGAGTCATTGTGATGAAATGCGTAGAGGAGAGGAGCGGGGAGGAGTTGGCGGCGGCGGAGGTTGGGCGGCGGCAGCCTGCGGCTAGGGC
TCCGCCCTGTTGTGCGGCGGGCGACTCGCGGCGGCGGCTGGGAACATTCGTGCTGCGGTGCGCGTGTTGTGAGGCTTGCTGTCGATTAAATTGGGCGCGT
GCCCCTGCAGGGACATCCCGGGTGGTGTAGGTCTGCAGAGGGTACGGATGGGTGGCGACGGTGCTTGGGCCGGATTGAGGGTTGGGTAAAGGCCTGCGAA
GCTTCAGTCGACGCGATTGTGGTGAGGGGGCTGGTGGTGGATCCCGCTTGTGGCGGTTATGCCCGGGCGCCAGTTTTGATACCGAGGGGGGACGTTGCAC
CGCTAGGGGCGTTAGCACGTGCCGGCGGGCAATTCCTCTTTTTGTCTTTGATGACGAGGATGTGAGTGGTGAATGCTTCTCCATGCTTGCGCATCGCATG
CGCCAGCATATAGATAGATAAAAAGTTGCACGACTGCGAGCGGTGGCGCAGCCCGTGCAATTGAGTAACGGTGCTGGCACATGGAAAGTGTCCATATGTT
CTTTGATTGCCGCGTGACCCCGCCACCGGTGATTCTTGCAAGCGCCTTATCCGGGCGCACATTCTGTAATTTCTGAGCCAGAATGATCAGGGTAGTGAGT
CCGGGTACAGGCTGCAGTGTCTGAGTTAAAGCTCGCAGGGTCAGGGTTACAGGGCAAAGGGTCAGGGGCAGGGGAGTAAGGGACCAGGGGTCAGGCGAGG
TTCAGCGGAGCTGGCTCGGGCTGTGTGAAGGTACAGGGGACGCTTAAGGTCAGGAGGCCAACAGTCAGAGTCAAGGGGGCCTCAGAGTCAGGGTGCCATC
AGCGTTATGTCTGCGAATGCAAGACAGGCGGCATTACGCGGTGCGTGCGGTCCAAGGGCTGTCAGGAACAGGCAATGCCCG

**Figure S11. *k6#* mutant complementation.** (a) genomic sequence scheme of the CP26 (Cre16.g673650) gene according to *Chlamydomonas reinhardti*i v 5.6 genome. 5' and 3' UTR are respectively reported in yellow and light blue, while CP26 gene is in green with introns represented as dotted lines. The putative promoter (1000 bp upstream) is represented with a gray arrow with red and blue regions referring to high complexity and un-sequenced regions. The putative terminator (300 bp downstream) is orange colored. (b) scheme of sequence used for complementation with CP29 upstream sequence. Putative CP29 promoter (1000 bp upstream) is represented with a blue arrow. Putative terminator (300 bp downstream) is orange colored (c) CP26 putative promoter region. High complexity and un-sequenced regions are in blue and red respectively. (d) Sequence of CP26 sequence used for complementation. Each part is colored as in (a), and introns are crossed out.

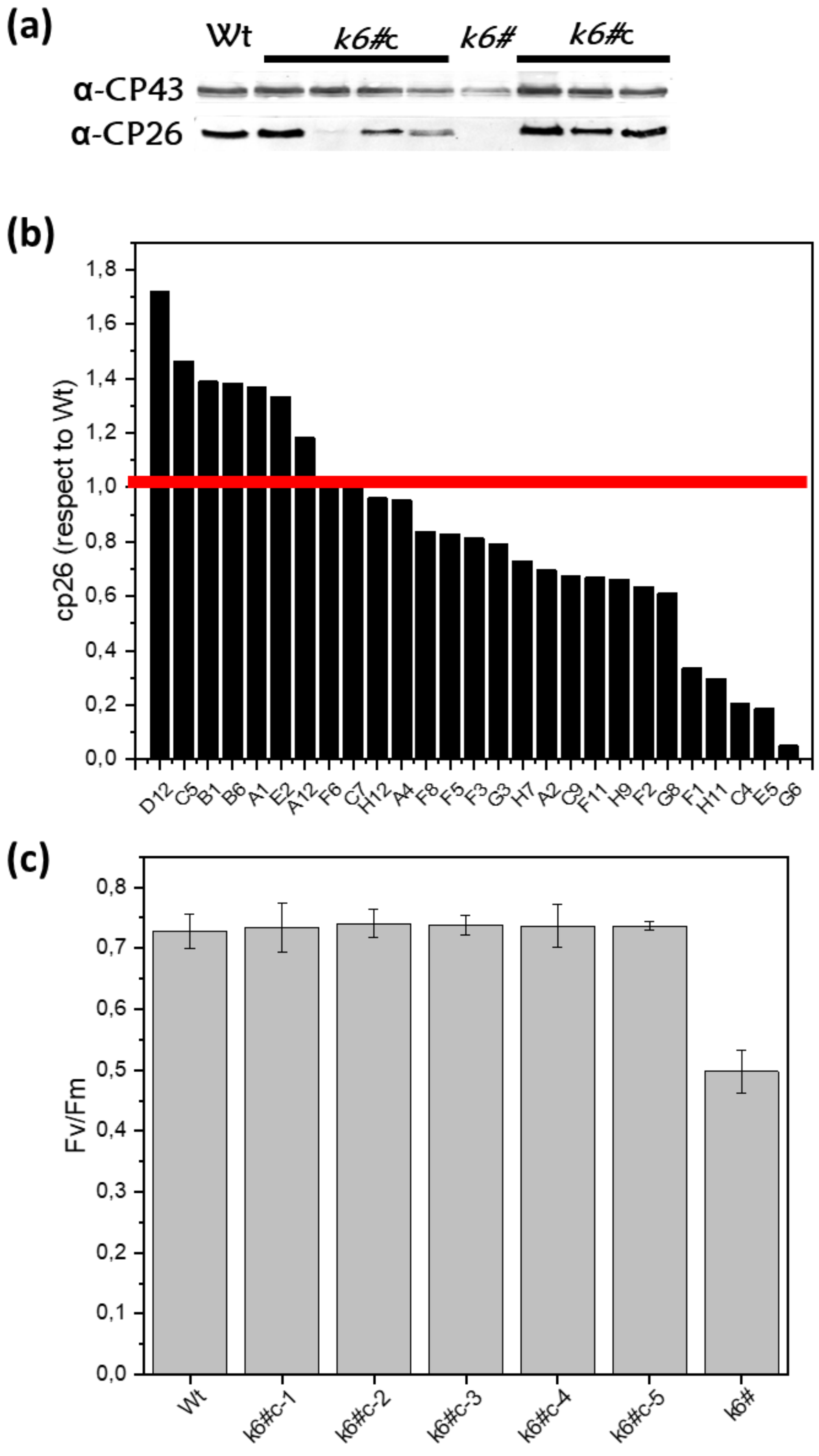


**Figure S12. *k6#* line complemented with CP26**. (a) example of immunotitration by western blot of the *k6#* mutant (line *k6#8*) complemented with CP26 (*k6#c*). Protein extract of *k6#c* lines were loaded on SDS-PAGE gel, transferred on nitrocellulose membrane, and blotted with specific antibodies against CP26 and CP43 (used to normalize loading differences). (b) CP26 amount in the different *k6#c* lines, data are normalized to the CP26/CP43 ratio of the wild type (Wt), indicated with the red line. (c) Fv/Fm of Wt, *k6#c* and *k6#* lines after two days in high light.

**Table S1. Sanger sequencing of the CP26 knock-out mutants.** The Indel value indicates the nucleotides inserted or deleted, compared to the sequence in wild type (Wt). The Hygromycin-resistance gene (HygR) cassette, which was co-transformed to help screening process, was confirmed that inserted forward (F) or reverse (R) direction in the genome. The original size of the HygR cassette is 1,624 bp and was included in the Indel size. The deleted sequences in the sgRNA target site were highlighted by gray colored line.

| | *CP26* sgRNA target site | Indel |
|---|---|---|
| Wt | CCCGAGGGCCTGCAGGC ------------- CAACGG | - |
| *k6#1* | CCCGAGGGCCTGCAGGC **(HygR (F))** ~~CAAC~~GG | +1975/-4 |
| *k6#2* | CCCGAGGGCC~~TGCAGGC~~ **(HygR (R))** ~~CA~~ACGG | +1258/-9 |
| *k6#3* | CCCGAGGGCCTGCAGGC **(HygR (R))** ~~C~~AACGG | +1340/-1 |
| *k6#4* | CCCGAGGGCCTGCAGGC **(HygR (F))** CAACGG | +1606 |
| *k6#5* | CCCGAGGGCCTGCAGGC **(HygR (R))** ~~CAA~~CGG | +1298/-3 |
| *k6#6* | CCCGAGGGCCTGCAGGC **(HygR (F))** CAACGG | +730 |
| *k6#7* | CCCGAGGGCCTGCAGGC **(HygR (F))** CAACGG | +1530 |
| *k6#8* | CCCGAGGGCCTGCAGGC **(HygR (F))** CAACGG | +1264 |
| *k6#9* | CCCGAGGGCCTGCAGG~~C~~ **(HygR (R))** ~~CA~~ACGG | +1249/-3 |
| *k6#10* | CCCGAGGGCCTGCAGGC **(HygR (F))** ~~CA~~ACGG | +1304/-2 |
| *k6#11* | CCCGAGGGCCTGCAGGC **(HygR (R))** ~~CA~~ACGG | +1512/-2 |

**Table S2. Fluorescence lifetimes of whole *Chlamydomonas reinhardtii* cells.** Fluorescence decay kinetics reported in Figure 3a and Figure S3b were fitted with 3-exponential function. The amplitudes (a1, a2, a3) and time constants (τ1, τ2, τ3) retrieved are reported with the average fluorescence lifetime (τ avg) calculated as (a1*τ1+a2*τ2+a3*τ3)/(a1+a2+a3). τ3 has been assumed constant to the value obtained by fitting Wt. The average value and standard deviations resulting from the fitting analysis are reported.

| Sample | $t_1$ (ps) | $a_1$ (%) | $t_2$ (ps) | $a_2$ (%) | $t_3$ (ps) | $a_3$ (%) | $t_{avg}$ (ps) |
|---|---|---|---|---|---|---|---|
| Wt | 96 ± 4 | 48.0 ± 1.1 | 497 ± 14 | 49.0 ± 1 | 2850 | 3.0 ± 0.3 | 375 ±11 |
| *k6#* | 99 ± 4 | 52.9 ± 1.3 | 504 ± 22 | 37.2 ± 1.1 | 2850 | 9.9 ± 0.4 | 522 ± 14 |
| *k9* | 97 ± 4 | 53.5 ± 1.2 | 534 ± 23 | 39.4 ± 1 | 2850 | 7.1 ± 0.5 | 465 ± 16 |
| *k69* | 103 ± 4 | 48.5 ± 1.4 | 491 ± 20 | 40.8 ± 1.2 | 2850 | 10.7 ± 0.4 | 555 ± 14 |

**Table S3 Light curve parameters and respiration rates.** Parameters extrapolated from oxygen light saturation curves shown in Figure 4d. Error bars are reported as standard deviation (n=3). *k6#* values that are significantly different (Student's t-test, $P < 0.05$) from wild type (Wt) are marked with *.

| | Wt | *k6#* |
|---|---|---|
| Respiration in the dark ($O_2$ nmol cell*$10^{-6}$ $min^{-1}$) | -2,05 ± 0,1 | -2,32 ± 0,67 |
| Pmax (O2 nmol ug $chl^{-1}$ $min^{-1}$) | 10,41 ± 1,17 | 10,5 ± 0,45 |
| Half-saturation intensity (µmol $m^{-2}$ $s^{-1}$) | 235 ± 13* | 320 ± 16* |
| slope of linear increase | 0,022 ± 0,001* | 0,018 ± 0,002* |
